# A feedback control mechanism governs the synthesis of lipid-linked precursors of the bacterial cell wall

**DOI:** 10.1101/2023.08.01.551478

**Authors:** Lindsey S. Marmont, Anna K. Orta, Robin A. Corey, David Sychantha, Ana Fernández Galliano, Yancheng E. Li, Becca W.A. Baileeves, Neil G. Greene, Phillip J. Stansfeld, William M. Clemons, Thomas G. Bernhardt

## Abstract

Many bacterial surface glycans such as the peptidoglycan (PG) cell wall, O-antigens, and capsules are built from monomeric units linked to a polyprenyl lipid carrier. How this limiting lipid carrier is effectively distributed among competing pathways has remained unclear for some time. Here, we describe the isolation and characterization of hyperactive variants of *Pseudomonas aeruginosa* MraY, the essential and conserved enzyme catalyzing the formation of the first lipid-linked PG precursor called lipid I. These variants result in the elevated production of the final PG precursor lipid II in cells and are hyperactive in a purified system. Amino acid substitutions within the activated MraY variants unexpectedly map to a cavity on the extracellular side of the dimer interface, far from the active site. Our structural evidence and molecular dynamics simulations suggest that the cavity is a binding site for lipid II molecules that have been transported to the outer leaflet of the membrane. Overall, our results support a model in which excess externalized lipid II allosterically inhibits MraY, providing a feedback mechanism to prevent the sequestration of lipid carrier in the PG biogenesis pathway. MraY belongs to the broadly distributed polyprenyl-phosphate N-acetylhexosamine 1-phosphate transferase (PNPT) superfamily of enzymes. We therefore propose that similar feedback mechanisms may be widely employed to coordinate precursor supply with demand by polymerases, thereby optimizing the partitioning of lipid carriers between competing glycan biogenesis pathways.

## MAIN TEXT

Bacterial cells surround themselves with a complex envelope that is essential for their integrity and shape. The envelopes of gram-negative (diderm) bacteria also serve as a formidable barrier against the entry of drug molecules, providing organisms like *Escherichia coli* and *Pseudomonas aeruginosa* with a relatively high intrinsic resistance to antibiotics^1, 2^. Understanding how these bacteria construct their envelope and regulate the assembly process therefore promises to aid in the identification of new vulnerabilities in surface biogenesis to target for antibiotic development.

The diderm envelope consists of two membranes: a cytoplasmic (inner) membrane and an asymmetric outer membrane (OM) with an inner leaflet of phospholipids and an outer leaflet composed of lipopolysaccharide (LPS)^2^. The LPS molecule consists of a Lipid A moiety, a core oligosaccharide, and a long polysaccharide chain called the O-antigen (O-Ag) that varies in composition between different strains and species^3^. Between the inner and outer membranes is the periplasmic space where the peptidoglycan (PG) cell wall is assembled. The PG layer is the essential stress bearing portion of the envelope that protects cells from osmotic lysis. It is constructed from glycan strands with repeating *N*-acetylglucosamine (GlcNAc) and *N*-acetylmuramic acid (MurNAc) sugars that are crosslinked by peptide stems attached to MurNAc forming the interconnected meshwork that encases the inner membrane^4^.

Surface glycans like PG and O-Ag are polymerized from monomeric building blocks attached to polyprenyl lipids via a pyrophosphate linkage. The lipid carrier is regenerated during polymerization, making it available for the continued production of monomer units to support synthesis of growing polymers^5, 6^. This synthetic strategy is conserved throughout biology with uses ranging from surface glycan biogenesis in microbes to the production of N-linked glycans in eukaryotic cells^7–10^. In any given organism, a common polyprenyl lipid carrier is used to produce multiple different glycans^7^. The concentration of these carriers is limiting^8^, suggesting that their utilization to produce monomer units for different pathways must be coordinated with the corresponding glycan polymerization process. Otherwise, excess accumulation of monomer units for one polymer will sequester the limiting carrier, indirectly inhibiting the production of other glycans that require the carrier for their biogenesis. Such precursor sequestration can have significant detrimental consequences for the cell envelope^8–11^. Despite the importance of efficient carrier utilization for the balanced synthesis of different surface glycans, the underlying mechanism has remained elusive.

The membrane-anchored precursor for PG biosynthesis is lipid II. Its synthesis begins in the cytoplasm, where multiple enzymes (MurA-F) sequentially assemble the activated sugar uridine diphosphate-MurNAc-pentapeptide (simplified as UM5)^4^. The phospho-MurNAc-pentapeptide moiety from this intermediate is then transferred to the lipid carrier undecaprenyl phosphate (C55P) at the inner face of the cytoplasmic membrane by the essential integral membrane enzyme MraY, generating the penultimate PG precursor, lipid I. The peripheral membrane enzyme MurG then transfers GlcNAc from UDP-GlcNAc to lipid I, forming the lipid II molecule, which contains the basic monomeric unit of PG linked to the C55 lipid by a pyrophosphate. Following its synthesis, lipid II is transported across the cytoplasmic membrane by the flippase MurJ where it can then be polymerized and crosslinked by PG synthases to form the cell wall matrix^12^.

There are two general types of PG synthases, and bacterial cells typically encode multiple members of each. The class A penicillin-binding proteins (aPBPs) are one type. They are single-pass membrane proteins with a large extracytoplasmic domain that possesses both PG glycosyltransferase (PGTase) activity to polymerize lipid II and transpeptidase (TPase) activity to crosslink the nascent glycans into the mature wall^13, 14^. The second type of synthase is formed by a complex between a SEDS (shape, elongation, division, and sporulation) family PGTase and a class B PBP (bPBP) with TPase activity. The SEDS-bPBP complexes form the essential PG synthases of the cell elongation and division machineries whereas the aPBPs are thought to fortify a foundational PG structure laid down by the morphogenic SEDS-bPBP systems^15–17^.

This investigation started with the study of *P. aeruginosa* mutants with a conditionally lethal defect in aPBP activity^18^. We isolated suppressors encoding an altered MraY enzyme with a T23P substitution [MraY(T23P)] that restored the growth of these cells in the non-permissive condition. Our characterization of this and other related MraY variants supports a model in which MraY is feedback inhibited by the accumulation of flipped lipid II, limiting the synthesis of PG precursors when their supply exceeds the synthetic capacity of PG synthases.

## RESULTS

### An MraY variant rescues a lethal aPBP synthase defect

*P. aeruginosa* produces two aPBPs, *^Pa^*PBP1a and *^Pa^*PBP1b, encoded by the *ponA* and *ponB* genes, respectively. These PG synthases require cognate OM lipoprotein activators to function properly^18, 19^. PBP1a is activated by *^Pa^*LpoA and *^Pa^*PBP1b is activated by *^Pa^*LpoP^18^ (**Fig. 1A**). A!i*ponB* !i*lpoA* mutant relies on an unactivated PBP1a enzyme for growth (**Fig. 1A**). We therefore refer to the strain as a PBP1a-only mutant for simplicity. Such mutants are viable on rich medium (lysogeny broth, LB), but were found to have severe growth defects on a defined minimal medium (Vogel-Bonner minimal medium, VBMM)^18^. Spontaneous suppressors supporting the growth of the PBP1a-only mutant on VBMM medium were isolated to uncover new insights into PG synthesis regulation. Several of these mutants were found to encode variants of *^Pa^*PBP1a, and we previously reported that they bypass the *^Pa^*LpoA requirement for *_Pa_*PBP1a function by activating the PG synthase^20^. Thus, the growth defect of the PBP1a-only strain on VBMM is caused by a deficit of aPBP activity. Here, we report the identification of a new class of suppressor with a mutation in *^Pa^mraY* encoding an enzyme variant with a T23P substitution [*^Pa^*MraY(T23P)] that alleviates the growth defect of the 1′*ponB* 1′*lpoA* strain.

**Figure 1.**
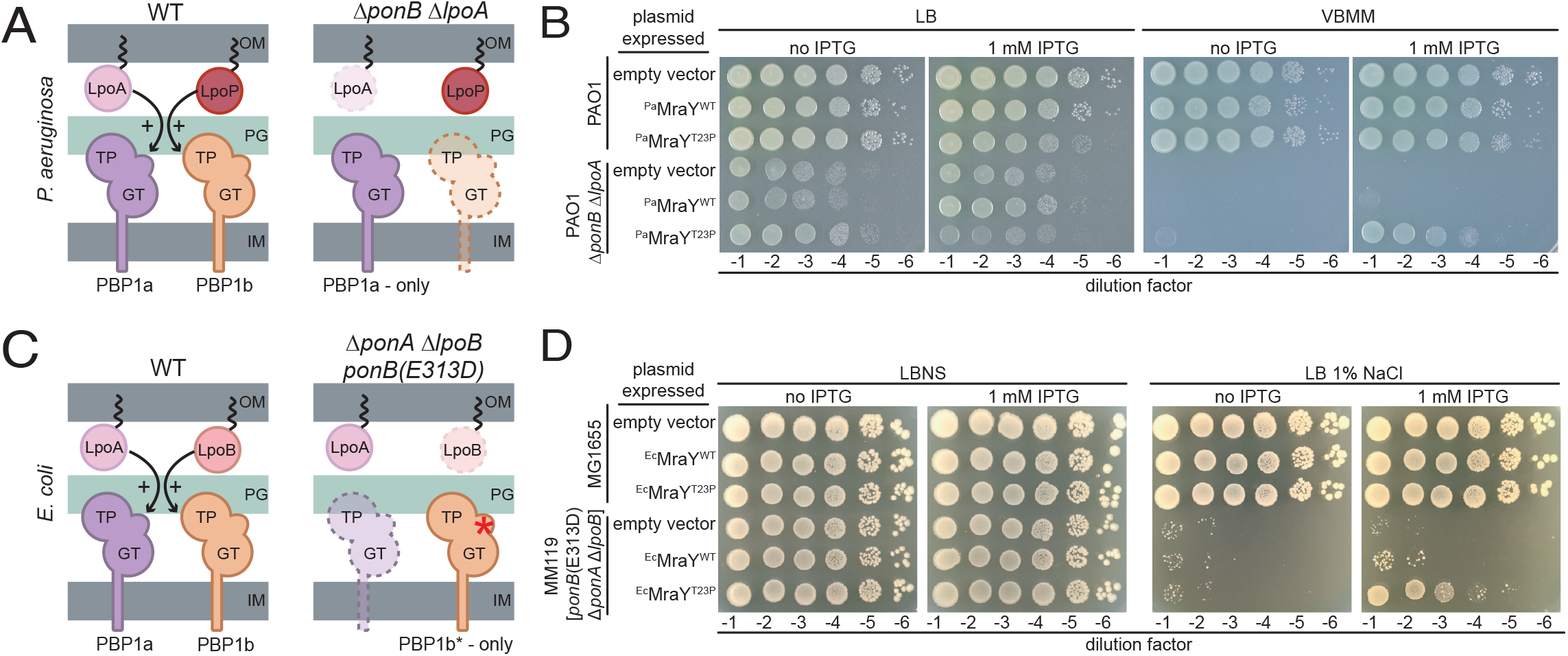
MraY(T23P) restores growth to strains defective in PG biosynthesis. Schematic representation of the aPBPs and their outer membrane lipoprotein activators in *Pseudomonas aeruginosa* (**A**) and *Escherichia coli* (**C**). Ten-fold serial dilutions of cells of the indicated *P. aeruginosa* (**B**) or *E. coli* (**D**) strains harboring expression plasmids producing the indicated MraY variant. Dilutions were plated on the indicated medium with or without IPTG to induce the production of MraY variants as indicated. Abbreviations: OM, outer membrane; PG, peptidoglycan; IM, inner membrane; GT, glycosyltransferase; TP, transpeptidase; LB, lysogeny broth; LBNS, LB with no added NaCl; VBMM, Vogel-Bonner minimal medium; IPTG, isopropyl-B-D-1-thiogalactopyranoside.

To confirm the suppression of the PBP1a-only growth defect by the MraY variant, *^Pa^mraY*(WT) or *^Pa^mraY*(T23P) were expressed from a multicopy plasmid under the control of an IPTG-inducible promoter in a wild-type *P. aeruginosa* (strain PAO1) or a 1′*ponB* 1′*lpoA* background **(Fig. 1B)**. Overexpression of *^Pa^mraY*(WT) in the wild-type PAO1 strain did not appreciably affect growth on either LB or VBMM nor did it rescue the lethal phenotype of the 1′*ponB* 1′*lpoA* mutant on VBMM **(Fig. 1B)**. Consistent with the results of the genetic selection, expression of *_Pa_mraY*(T23P) restored growth of the PBP1a-only mutant on VBMM with as little as 25 µM of inducer promoting significant growth on these non-permissive conditions **(Fig. 1B and Fig. S1)**. Notably, in addition to rescuing growth of the mutant on VBMM, overexpression of *_Pa_mraY*(T23P) caused a mild growth defect in both PAO1 and 1′*ponB* 1′*lpoA* backgrounds when cells were grown on rich medium **(Fig. 1B and Fig. S1)** (see below). Substitution of the catalytic residue D267 with Ala in the active site of *^Pa^*MraY(T23P) eliminated the toxicity of the variant when it was overproduced in PAO1 cells on LB and greatly reduced the suppression activity in 1′*ponB* 1′*lpoA* cells on VBMM **(Fig. S1)**. Furthermore, VSVG-tagged derivatives of *_Pa_*MraY(WT) and *^Pa^*MraY(T23P) were found to accumulate to similar levels in cells by immunoblot analysis with the tagged *^Pa^*MraY(T23P) variant promoting better growth of 1′*ponB* 1′*lpoA* cells on VBMM than the tagged wild-type protein **(Fig. S2)**. Thus, the suppression activity of the *^Pa^*MraY(T23P) variant is not due to increased accumulation of the enzyme. Rather, the results suggest that the T23P change alters MraY activity to promote the growth of the aPBP deficient strain on VBMM and impair growth of both mutant and wild-type strains on LB when it is overexpressed.

*E. coli* also encodes aPBPs, *^Ec^*PBP1a and *^Ec^*PBP1b, controlled by OM lipoprotein activators *_Ec_*LpoA and *^Ec^*LpoB, respectively (**Fig. 1C**)^19, 21^. We previously described an *E. coli* strain lacking *^Ec^*PBP1a and *^Ec^*LpoB that relies on a LpoB-bypass variant of *^Ec^*PBP1b [*^Ec^*PBP1b(E313D)] as its only aPBP (**Fig. 1C**)^22^. Like the *P. aeruginosa* 1′*ponB* 1′*lpoA* strain, this *E. coli* mutant has a conditional growth defect caused by a deficit in aPBP activity. It grows on LB without added NaCl (LBNS) but is inviable on LB with 1% NaCl. Overproduction of *E. coli* MraY(T23P) [*^Ec^*MraY(T23P)] but not wild-type *^Ec^*MraY suppressed the growth defect of this aPBP-deficient *E. coli* strain on LB 1% NaCl (**Fig. 1D**). Therefore, an MraY(T23P) variant suppresses an aPBP defect in two distantly related gram-negative bacteria, suggesting that its properties are conserved.

### MraY(T23P) is activated and increases lipid II accumulation in cells

MraY uses UM5 and C55P to form the first lipid-linked PG precursor lipid I, which is then converted to the final precursor lipid II by MurG. We reasoned that MraY(T23P) might overcome the aPBP-deficiency in mutants of *P. aeruginosa* and *E. coli* by increasing the concentration of the synthase substrate lipid II to compensate for the poorly activated aPBP in these cells. To investigate this possibility, we measured the concentration of lipid II in *P. aeruginosa* and *E. coli* cells overproducing MraY(WT) or MraY(T23P). Exponentially growing cultures were normalized by optical density, and the cells were harvested and extracted for lipid-linked PG precursors (**Fig. 2A**). The extract was subjected to acid hydrolysis to release the disaccharide-pentapeptide from undecaprenyl-pyrophosphate (C55PP), and the soluble disaccharide-pentapeptide was subsequently detected by liquid chromatography/mass spectrometry (LCMS) as a measure of lipid II concentration **(Fig. 2B-E)**. In both the wild-type and aPBP deficient mutant backgrounds, MraY(WT) overproduction led to an approximately twofold increase in lipid II levels relative to an empty vector control **(Fig. 2C and 2E)**. The increase was another twofold higher for cells overproducing MraY(T23P) **(Fig. 2C and 2E)**. We observed similar trends monitoring lipid I levels, but the increase in lipid I levels in cells producing MraY(T23P) relative to MraY(WT) was not nearly as pronounced compared to the change in lipid II levels (**Fig. S3**). These results suggest that the altered MraY enzyme is more active than wild-type and that the ability to promote the accumulation of higher lipid II levels indeed underlies the suppression of aPBP defects.

**Figure 2.**
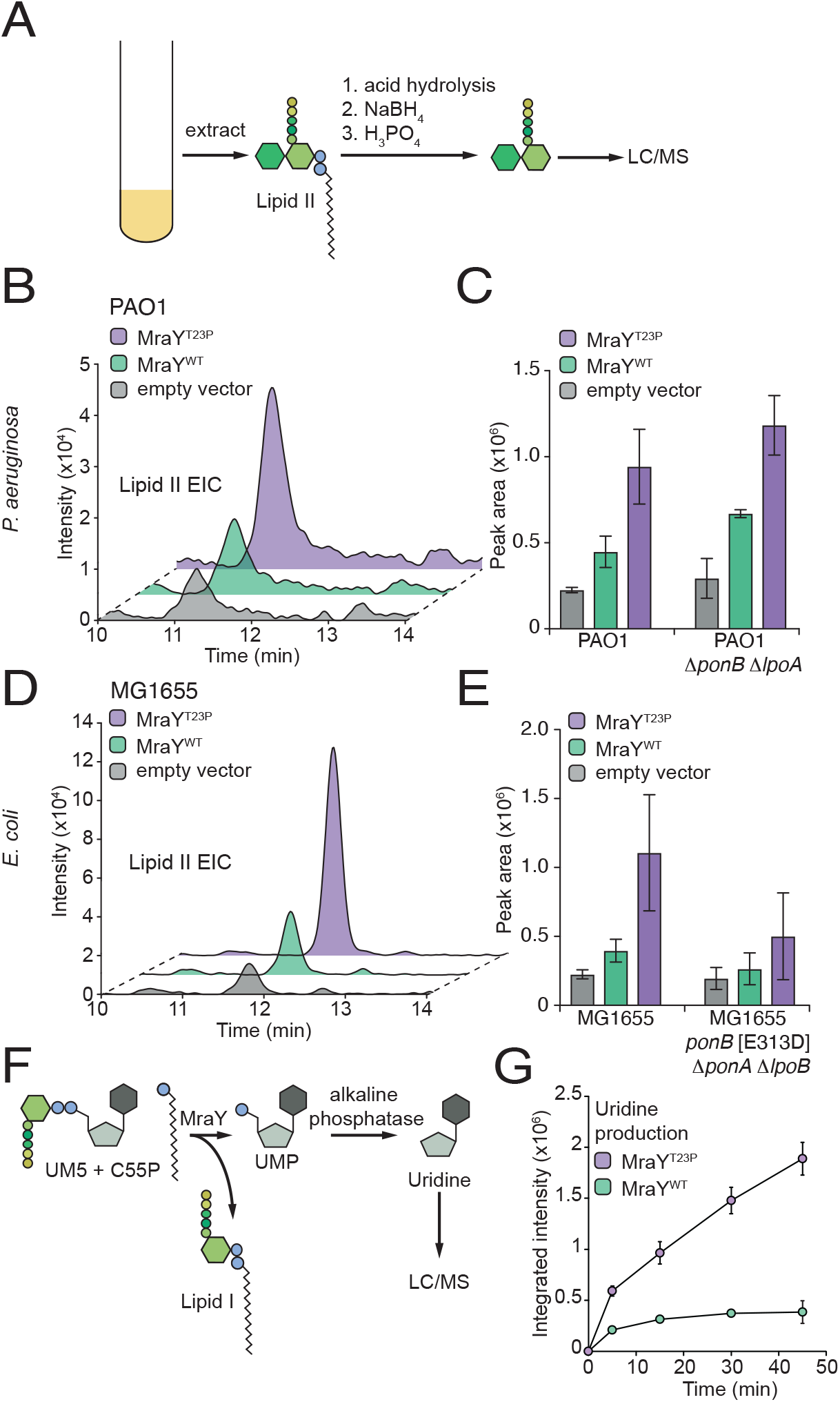
Cells expressing MraY(T23P) accumulate lipid II. (**A**) Schematic representation of the method used to isolate and analyze lipid II from bacterial cells. (**B-D**) Representative extracted ion chromatograms of lipid II (EIC) and quantification of EICs for *P. aeruginosa* (**B,C**) or *E. coli* (**D,E**) strains expressing the indicated MraY variant. Three independent replicates of the extractions were performed and lipid II levels quantified using the area of the peak from the extracted ion chromatogram using the Agilent software. Error bars represent the standard deviation. For MraY(T23P) vs MraY(WT) in PAO1 P<0.05 PAO1 1′*ponB* 1′*lpoA* P<0.01, in PAO1, MG1655 P<0.05, MG1655 1′*ponA* 1′*lpoB ponB*[E313D], not significant. (**F**). Schematic representation of the MraY enzyme assay. (**G**) Representative time course showing the production of uridine in assays containing purified MraY or MraY(T23P) as indicated. The assay was repeated at least twice with two independent preparations of protein. Error bars represent standard deviation of reactions performed in duplicate. Abbreviations: NaBH_4_, sodium borohydride; H_3_PO_4_, phosphoric acid; LC/MS, liquid chromatography mass spectrometry; UM5, UDP-MurNAc-pentapeptide; C55P, undecaprenylphosphate.

To assess the effect of the T23P substitution on MraY activity directly, FLAG-tagged derivatives of *^Pa^*MraY(WT) and *^Pa^*MraY(T23P) were heterologously expressed in *E. coli* and affinity purified for biochemical assays. The reaction was followed by monitoring the production of uridine derived from alkaline phosphatase treatment of the UMP product (**Fig. 2F**). Using this assay, the *^Pa^*MraY(T23P) variant was found to be significantly more active than *_Pa_*MraY(WT). At the conclusion of the time-course, approximately five times more uridine was detected in reactions containing *^Pa^*MraY(T23P) than those with *^Pa^*MraY(WT) (**Fig. 2G**). We conclude that the T23P substitution generates a hyperactive MraY, leading to elevated lipid II production in cells.

### Misregulated MraY disrupts O-antigen synthesis

The results thus far suggest that *^Pa^*MraY(T23P) makes more lipid-linked PG precursors than normal, leading to their hyperaccumulation. In the PBP1a-only *P. aeruginosa* strain, this increase in substrate supply suppresses the lethal aPBP deficiency. We wondered whether excess lipid II production and the resulting sequestration of C55P in this building block might also indirectly impede the synthesis of other surface glycans built on the lipid carrier like O-Ag. A clue that this was the case came from the growth defect on LB medium of the wild-type PAO1 strain caused by *^Pa^*MraY(T23P) but not *^Pa^*MraY(WT) overproduction **(Fig. 1B and Fig. S1)**. Notably, this strain produces R2-pyocin, a lethal phage tail-like bacteriocin that uses a receptor located within the LPS core to engage target cells^23, 24^. *P. aeruginosa* is resistant to killing by its own R2-pyocin because it decorates its LPS with O-Ag that masks the R2-pyocin receptor. Defects in the O-Ag synthesis pathway therefore result in susceptibility to R2-pyocin self-killing^25^. The connection between O-Ag and R2-pyocin activity suggested to us that the growth phenotype induced by *^Pa^*MraY(T23P) overproduction on LB medium may be caused by a decrease in O-Ag production and increased R2-pyocin self-intoxication. To test this possibility, we examined the effect of *^Pa^*MraY(T23P) overproduction in a strain deleted for the R2-pyocin gene cluster (*PA0615-PA0628*). Strikingly, unlike wild-type cells, the mutant incapable of making R2-pyocin was largely unaffected by the overproduction of *^Pa^*MraY(T23P) **(Fig. S4A)**, indicating that the growth defect caused by the altered enzyme was largely due to R2-pyocin killing. This result suggested that O-Ag synthesis is reduced when lipid II synthesis is hyperactivated in cells producing *^Pa^*MraY(T23P). Analysis of the LPS produced by these cells confirmed that they indeed have reduced levels of O-Ag. They made approximately 30% less O-Ag compared to cells expressing *^Pa^*MraY(WT) (**Fig. S4B-C**). These results suggest that *_Pa_*MraY(T23P) may be insensitive to a regulatory mechanism limiting the steady-state accumulation of lipid-linked PG precursors to prevent the impairment of competing pathways utilizing the C55P carrier.

### The extracytoplasmic side of the MraY dimer interface may be a regulatory site

MraY is a polytopic membrane protein with ten transmembrane helices and an N-out, C-out topology^26^. The structure of the enzyme from *Aquifex aeolicus* revealed that it forms a dimer with most of the monomer-monomer contacts made between the N-and C-terminal helices^26^. Notably, the T23 residue lies near the dimer interface on the extracytoplasmic side of MraY. We therefore wondered whether other substitutions in this area might also activate the enzyme. To test this possibility, a mutagenized copy of *^Pa^mraY* under the control of an IPTG inducible promoter was transformed into the 1′*ponB* 1′*lpoA P. aeruginosa* strain. The resulting transformants were then selected on VBMM in the presence of IPTG to identify MraY variants that rescue the aPBP deficiency. Twenty-one suppressing clones were isolated that each contained a single point mutation in the plasmid-borne copy of *mraY* (**Fig. 3A**). The positions of these substitutions were mapped onto a model of the *^Pa^*MraY structure generated using AlphaFold^27, 28^. Strikingly, all changes were located proximal to the dimer interface, with a majority positioned on the extracytoplasmic side of the protein far from the active site, which is located on the cytoplasmic side of the enzyme (**Fig. 3B-C, Table S1**). Overall, our genetic and biochemical results implicate the extracytoplasmic region of MraY near the dimer interface as a potential regulatory site for the enzyme.

**Figure 3.**
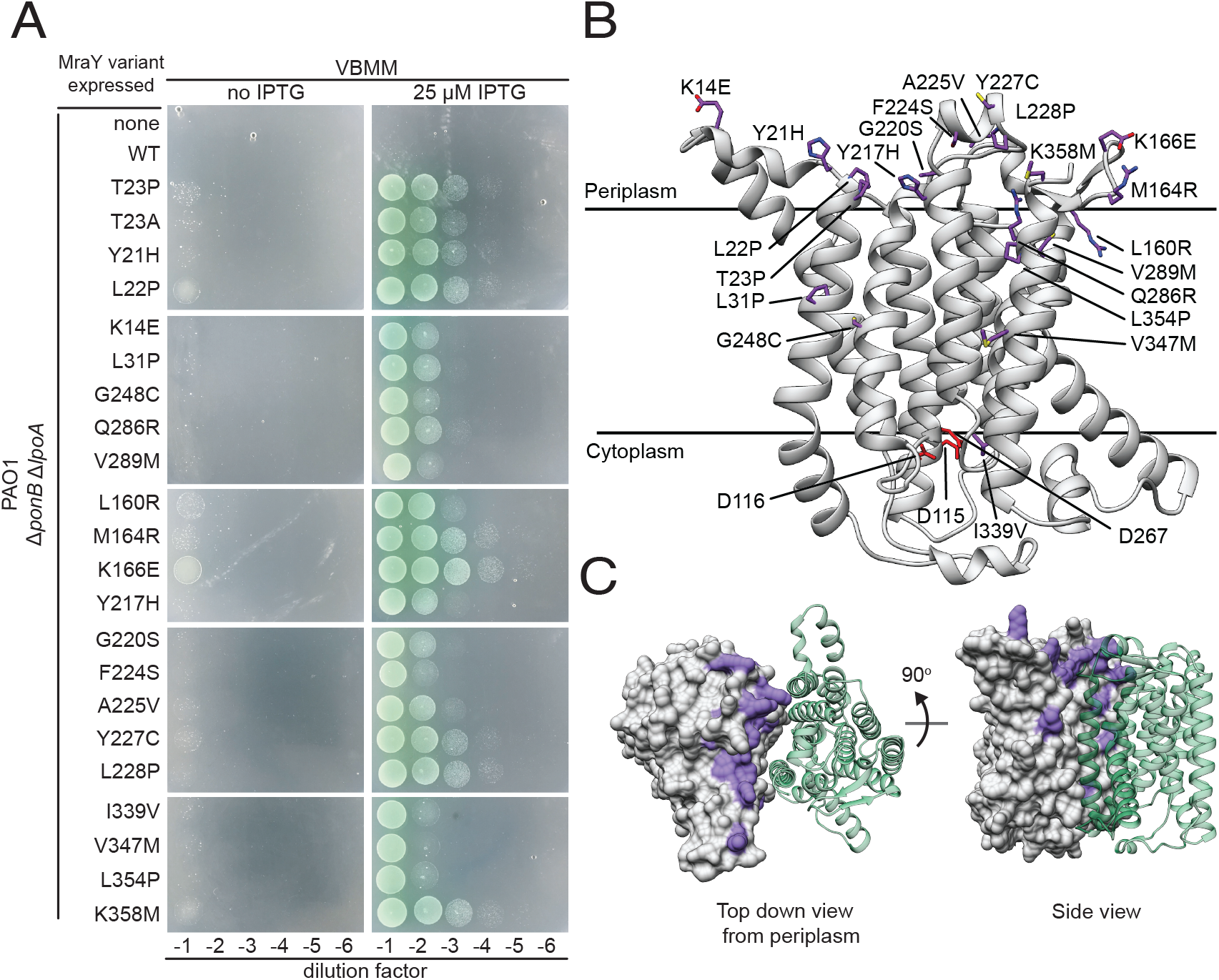
Amino acid substitutions in hyperactive MraY variants localize to the extracytoplasmic surface of the dimer interface. (**A**) Ten-fold serial dilutions of *P. aeruginosa* 1′*ponB* 1′*lpoA* cells harboring expression plasmids producing the indicated MraY variant were plated on VBMM with or without IPTG to induce the MraY variants as indicated. (**B**) Structural model of *P. aeruginosa* MraY created using AlphaFold^28^ in cartoon viewed from the plane of the membrane. Residues altered in hyperactive variants tested in (**A**) are shown in stick representation (purple), while those residues previously implicated in catalysis are shown in red. (**C**) Structural model of the MraY dimer created using AlphaFold^28^. Surface representation of one protomer is shown in grey with the residues altered in hyperactive variants colored in purple. The other protomer is shown in cartoon representation (green) for simplicity. Left, the periplasmic view of the dimer; Right, view from the plane of the membrane.

### A potential binding site for flipped lipid II within the MraY dimer interface

Both the *A. aeolicus* and *Enterocloster boltae* MraY crystal structures revealed the presence of a cavity located at the dimer interface that is lined by hydrophobic residues^26, 29^. The authors concluded that electron density within this tunnel could accommodate a cylindrical molecule that is too long to be detergent from the sample preparation^26^. Instead, they suggested that this electron density could accommodate one or more lipid molecules. Although it has been speculated to be C55P^26^, the identity of the lipid has remained unclear. Additionally, a recent study identified lipid molecules co-purifying with MraY using native mass-spectrometry^30^. The most abundant species were the C55P substrate and lipid I product, but peaks corresponding to C55PP, cardiolipin, and lipid II were also detected^30^. Thus, MraY likely binds a lipid molecule within the dimer interface near residues we have implicated in controlling the activity of the enzyme.

Clues to the potential identity of the lipid bound at the MraY dimer interface came from structural analysis of *^Ec^*MraY in complex with a phage-encoded inhibitor (protein E) and the *E. coli* chaperone SlyD (the YES complex)^31^. The cryo-EM structure of the YES complex containing wild-type *^Ec^*MraY was recently reported^31^, and these methodologies were used to obtain the structure of *^Ec^*MraY(T23P) within the same complex (**Fig. S5**, **Table S2**). In both cases, electron density was observed at the MraY dimer interface. Focused refinement of MraY alone in the *^Ec^*MraY(T23P) complex significantly improved the potential lipid density at the MraY dimer interface **(Fig. 4A-B)**. As in previous *A. aeolicus* and *E. boltae* MraY structures, this electron density fills the hydrophobic cavity found at the MraY dimer interface. However, we uniquely observed this electron density extending into the periplasmic space above the MraY molecules where the environment is more hydrophilic (**Fig. 4A-B**). Although structural refinement alone could not conclusively identify the lipid within the dimer, the size of the electron density extending into the periplasmic space is consistent with a large head-group such as the disaccharide-pentapeptide found on lipid II.

**Figure 4.**
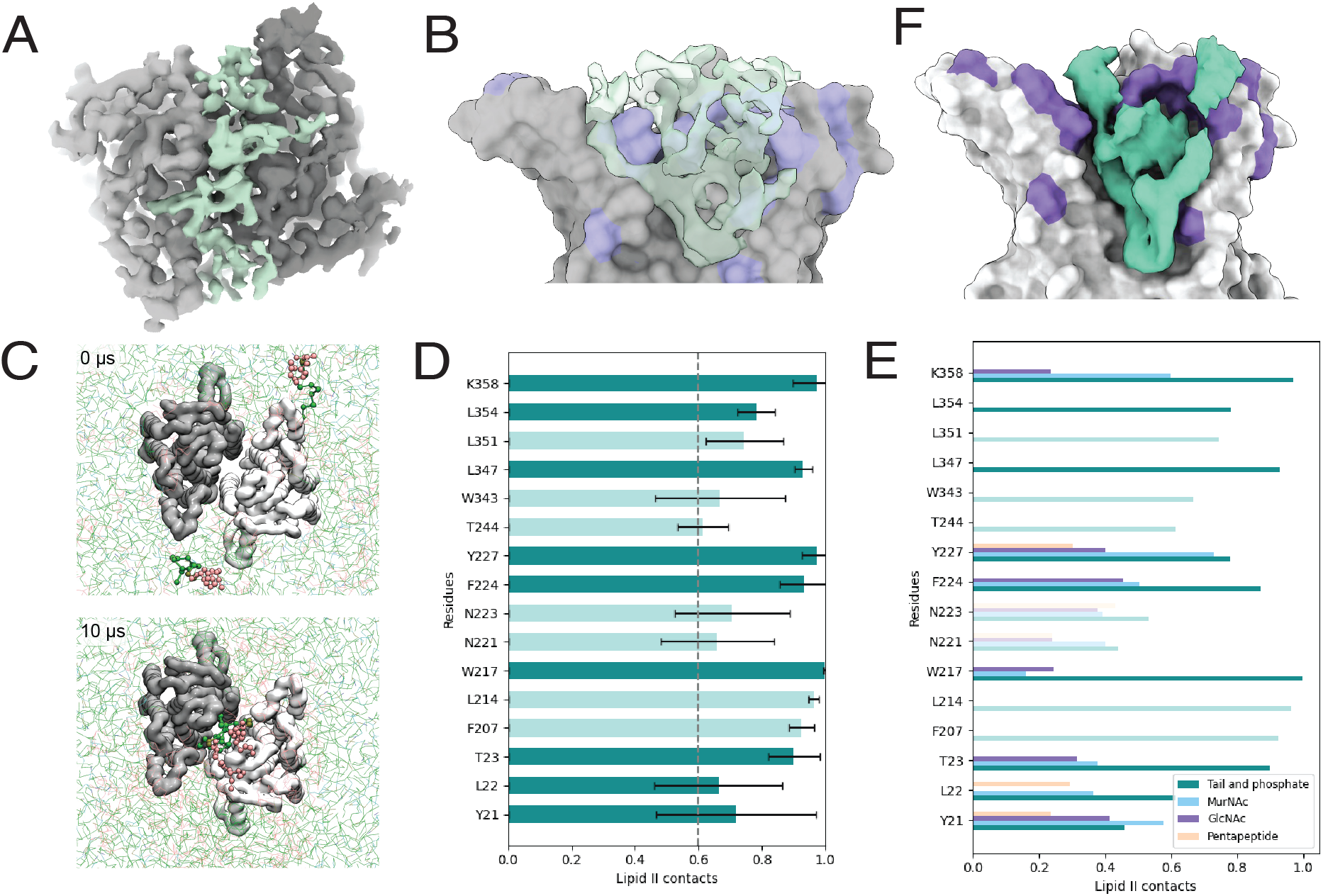
Identification of a potential lipid II binding site in MraY. (**A**) Periplasmic view of the cryo-EM structure of the *^Ec^*MraY(T23P) dimer within the YES complex shown in surface representation, with the unmodeled electron density shown (green). (**B**) As in *A*, membrane view of the electron density within the dimer interface of the *^Ec^*MraY dimer with the foreground MraY removed. (**C**) Top view of the MraY dimer in a mixed CG membrane. Two lipid II molecules (highlighted as green, gold and pink spheres) freely enter the MraY cavity during unbiased MD simulations. In 8/9 repeats, 2 or 3 lipid I or II molecules bind the cavity. In the last repeat, one lipid II and one C55P molecule bind. (**D**) Lipid II contacts with MraY residues that interact with lipid II for over 60% of atomistic MD simulations. Error bars represent standard error from 5 repeats. Darker green bars represent residues altered in hyperactive variants. (**E**) Lipid II contacts with MraY residues by part of lipid II that is interacting (tail & phosphate, MurNAc, GlcNAc or pentapeptide). Residues shown are same as those in Fig 4D. Darker bars represent residues altered in hyperactive variants. (**F**) Average density of lipid II molecules (green) from atomistic MD simulations of MraY (grey) bound to lipid II. Shown as inside view of dimer interface, where only one monomer of MraY is shown and residues altered in hyperactive variants are colored in purple.

To assess whether a lipid II molecule could enter the hydrophobic cavity of the MraY dimer, we used molecular dynamics (MD) simulations. In the first set of simulations, we used the structure of the *E. coli* MraY dimer from the YES complex (PDBID 8G01)^31^ embedded in a lipid bilayer [palmitoyl-2-oleoyl-*sn-*glycero-3-phosphoethanolamine (POPE), palmitoyl-2-oleoyl-*sn*-glycero-3-phosphoglycerol (POPG) and cardiolipin (CDL)] containing C55P, C55PP, lipid I, or lipid II with hydrophilic head-groups oriented towards what would be the periplasmic side of the membrane. Using course-grained MD simulations we observed that in almost all runs, lipid I and lipid II molecules spontaneously entered the central cavity, where typically two molecules would occupy the cavity (**Fig. 4C**, **Movie S1**). POPE and POPG never entered the cavity, while C55P and C55PP would occasionally enter the cavity at approximately 4% and 10% of the simulation time (**Movie S2**). Lipid I and Lipid II would remain stably bound for at least several µs (**Movie S1**), reflected by the K_off_ values for lipid I and lipid II of 0.218 µs^-1^ and 0.206 µs^-1^, respectively. In similar experiments where lipid I and lipid II are omitted, a single CDL molecule or C55P molecule will enter the cavity (**Movie S2**), although in the majority of simulations, no lipid enters the cavity. These were much shorter lasting interactions, with K_off_ values of 3.976 µs^-1^ for C55P and 1.297 µs^-1^ for CDL. Although lipid I and lipid II have similar K_off_ values in these simulations, lipid I is not found in the periplasmic leaflet of the inner membrane. Therefore the simulations with lipid I are not likely to reflect a physiologically relevant binding event. Instead, lipid II is the best candidate for the native ligand due to its strong and long-lasting interaction. Notably, the bound lipid II molecules in the simulations make extensive contacts with the MraY dimer, with many residues contacting the bound lipids for 100% of the MD simulations (**Fig 4D, Fig. S6**). These residues include several that were identified in the mutational analysis as being hyperactive (**Fig. 3A-B**). To investigate the interaction in more detail, a pose of the *E. coli* MraY dimer with two bound lipid II molecules was converted to an atomistic description for further MD analysis. The data show that the lipid II molecules are stable in the central cavity with the isoprenyl chains adopting a curved orientation. The result predicts contacts between the MurNAc sugar and MraY that include several residues where substitutions were identified in our screen (Y21, L22, T23, W217, F224, Y227, and K358) (**Fig. 4E-F**). Together, these data indicate that C55P-linked lipids can spontaneously enter a previously empty MraY dimer interface cavity and that externalized lipid II is likely to be the ligand bound in the potential regulatory site identified in the genetic and biochemical analyses.

## DISCUSSION

Bacterial surfaces contain multiple types of glycan and other polymers that are required for cellular integrity and/or barrier function. Although most of the proteins involved in the synthesis of major surface components are known, how the biogenesis of these molecules is regulated to efficiently distribute shared precursors like the C55P lipid carrier among competing synthesis pathways remains poorly understood. In this report, we uncover a mechanism governing the activity of MraY, the essential enzyme catalyzing the first membrane step in the PG synthesis pathway in which C55P is consumed to form lipid-linked PG precursors. This regulation is likely to play an important role in the efficient distribution of C55P among glycan biogenesis pathways that utilize the limiting carrier.

The first clue that MraY is regulated came from the discovery that an *mraY(T23P)* mutant can suppress an aPBP deficiency in both *P. aeruginosa* and *E. coli*. The aPBP deficient strains encode a single aPBP lacking its required activator. Prior work with these strains suggests that their conditionally lethal growth phenotypes are caused by poor PG synthesis efficiency resulting from the synthase having a reduced affinity for lipid II in the absence of its activator^20^. Accordingly, we infer that MraY(T23P) suppresses this problem by raising the steady state level of lipid II to overcome the substrate binding limitations of the unactivated aPBP. The ability of the altered MraY to increase lipid II levels indicates a role for the enzyme in regulating the maximum level of lipid II in cells. We propose that this control is mediated via feedback inhibition of MraY by externalized lipid II (**Fig. 5**).

**Figure 5.**
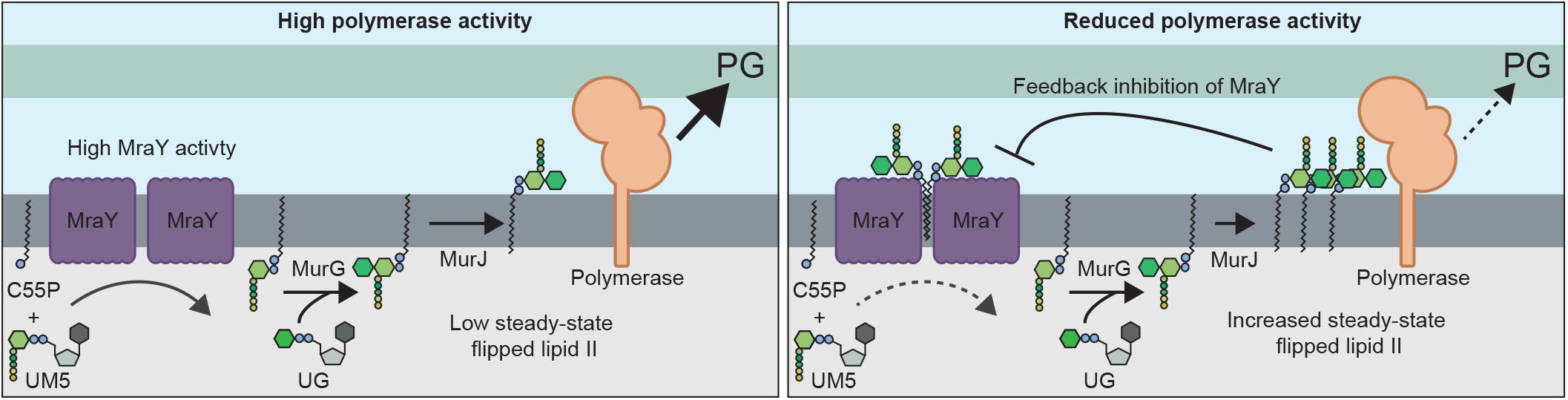
Model for feedback regulation of MraY by flipped lipid II. Shown are schematics summarizing the model for MraY regulation. Left: When PG polymerase activity is high, flipped lipid II is consumed at a rate proportional to its production such that steady-state levels of the precursor remains low and MraY activity is unimpeded. Right: When PG polymerase activity is reduced due to changes in growth conditions or other perturbations, lipid II will be produced faster than it is consumed, resulting in the accumulation of elevated levels of flipped lipid II. Higher levels of the precursor promote its binding to MraY dimers, reducing their activity in order to bring lipid II supply back in balance with demand by the polymerases. See text for details. Abbreviations: C55P, undecaprenylphosphate; UM5, UDP-MurNAc-pentapeptide; UG, UDP-GlcNAc; PG, peptidoglycan.

In support of the feedback inhibition model, the biochemical results with purified enzymes indicate that the observed regulation is intrinsic to MraY and does not require additional proteins. The MraY(T23P) variant, which is apparently less sensitive to regulatory control, showed much greater activity *in vitro* than the wild-type enzyme. At first glance, this result may seem incompatible with the proposed feedback control given that the product of the reaction is lipid I with its head-group in the cytoplasm, not externalized lipid II. However, because the reactions are performed in detergent, the lipid I formed in the reaction is likely capable of reorienting in the micelles to mimic a periplasmic orientation. Although externalized lipid I is not observed *in vivo*, the MD simulations predict that both flipped lipid I and lipid II are capable of binding at the MraY dimer interface. It is therefore reasonable to interpret the biochemical results in the context of a feedback inhibition model with MraY(WT) activity leveling off early in the time course due to feedback control. By contrast, we infer that MraY(T23P), with its substitution in the proposed binding site for flipped lipid II, is insensitive to feedback control and therefore displays robust activity in the assay. Another factor that is likely to contribute to the biochemical results is the co-purification of lipid II with the purified enzymes, which according to the model would be expected to further reduce the activity of MraY(WT) relative to MraY(T23P). Importantly, the activity for the wild-type enzyme was already so low that it was not possible to directly test for feedback inhibition via the addition of purified lipid II to the enzyme. Nevertheless, based on the logic above and the totality of the results presented, feedback inhibition of MraY by flipped lipid II provides one of the simplest and most straightforward explanations for our findings.

Although additional experiments are required to further investigate the possible feedback regulation of MraY, it is a compelling model because it suggests a mechanism by which cells can balance the supply of flipped lipid II precursor with the activity of the PG synthases that use it to build the cell wall (**Fig. 5**). We propose that when PG synthases are highly active, the steady state level of lipid II remains low such that MraY is functioning near its maximum activity to continue supplying lipid-linked PG precursors (**Fig. 5, left panel**). However, when the supply of lipid II exceeds the capacity of the PG synthases to use it, either transiently or due to a change in growth conditions, the steady state level of lipid II will rise such that it begins binding MraY dimers to inhibit their activity and reduce flux through the lipid stages of PG precursor production until supply more closely matches demand (**Fig. 5, right panel**). Such feedback control would prevent excess C55P from being sequestered in PG precursors when they are not needed, making more of the lipid carrier available to other glycan synthesis pathways for their efficient operation. Accordingly, *P. aeruginosa* cells with an activated MraY variant, which is presumably less sensitive to feedback control, display reduced ability to make O-Ag, rendering them susceptible to self-intoxication by their encoded pyocins (**Fig. S4B, C**).

The location of the amino acid substitutions in MraY that suppress aPBP defects combined with the structural and MD analysis suggest a mechanism by which the enzyme may be regulated by lipid II binding. Many of the MraY substitutions that overcome the PG synthesis defects of the PBP1a only strain localize to the extracytoplasmic surface of the protein distal to, and on the other side of the membrane from, the active site. These changes flank the opening of a deep hydrophobic pocket at the MraY dimer interface. In the cryo-EM structure of MraY within the YES complex^31^, we observe an MraY dimer with electron density at this interface as observed in prior X-ray crystal structures^26, 29^. However, in our structure, this density not only fills the pocket but also extends into the extracytoplasmic opening. This density in the extracytoplasmic space is large enough to correspond to a head-group of flipped lipid II. Accordingly, MD simulations indicate the capacity of MraY dimers to bind two molecules of flipped lipid II with contacts between the protein and the MurNAc sugar that likely provide specificity for externalized lipid II binding over C55PP or C55P. Notably, the head-groups of the lipid II binding substrates remain relatively flexible in the simulations (**Movie S1, Fig. S7**), which likely accounts for our inability to further refine the structure of the bound molecules by cryo-EM.

The MD simulations predict conformational changes in the MraY dimers associated with lipid II binding that increase the distance between the 6^th^ transmembrane helix (TM6) of each monomer in the dimeric structure and alter the position of the 9^th^ transmembrane helix (TM9) (**Fig. S8A-D**). Similarly, the distance between a periplasmic helix (residues 221-228) from each monomer is also increased (**Fig. S8C-F**). These changes are reminiscent of the conformational difference between MraY in the YES complex relative to the free MraY structure from *A. aeolicus*^26^. When the structures are aligned on one monomer, the second monomer in the YES complex^31^ is tilted relative to its partner in the *A. aeolicus* dimer^26^ resulting in the opening the periplasmic cavity and tightening the interface at the cytoplasmic side of the enzyme where the active site is located (**Fig. S9**). Because MraY in the YES complex is inhibited by the phage lysis protein, this opened conformation likely represents the inhibited state. The similarities between the conformational changes in MraY observed in the YES complex and upon lipid II binding in the MD analysis indicate that it is feasible for lipid II binding on the periplasmic side of the enzyme to be communicated to the active site via an alteration of the dimer interface. Accordingly, an increased mobility of TM9 on the cytoplasmic-face is also observed in the MD analysis when lipid II is bound (**Fig. S8B**). How the T23P and other changes that presumably activate MraY by reducing the sensitivity of the enzyme to inhibition by lipid II are not yet clear.

However, electron density corresponding to the lipid is still observed at the dimer interface between MraY(T23P) protomers in the variant YES complex. Although this result may be affected by the enzyme being stuck in an inhibited state by the phage inhibitor, it suggests that T23P and other changes in MraY may affect the conformational response of the enzyme to lipid II binding rather than the binding event itself. Consistent with this possibility, tyrosine at position 21 has an altered conformation in the MraY(T23P) structure in which its hydroxyl group forms a hydrogen bond network with Y227 and K358 on the opposing monomer (**Fig. S10**). Substitutions within these residues were also identified in the screen for hyperactive MraY enzymes, and Y227 is in the periplasmic helix that was found to be altered in the MD analysis upon lipid II binding. Thus, alterations affecting interactions in this region may be responsible for the regulation of MraY activity and its potential modulation by lipid II binding.

MraY belongs to the polyprenyl-phosphate N-acetylhexosamine 1-phosphate transferase (PNPT) superfamily of proteins that are found in all domains of life. The superfamily includes enzymes that initiate the lipid-linked stages of many glycan polymers including O-antigens, capsules, and teichoic acids in bacteria. A well-studied example outside of bacteria is the GlcNAc-1-P-transferase (GPT) that catalyzes the first step of N-linked protein glycosylation in eukaryotes by conjugating GlcNAc to the lipid carrier dolichol phosphate (DolP) to form Dol-PP-GlcNAc^11^. In each synthesis pathway, the final lipid-linked precursor for each glycan is built on a lipid carrier that must be shared with other pathways. It would therefore not be surprising if externalized versions of the final lipid-linked precursors of many different glycan biogenesis pathways exerted feedback control on the PNPT superfamily member that initiates precursor synthesis. Such a broad utilization of this feedback regulation would provide a mechanism to efficiently distribute limiting lipid carrier molecules between competing glycan synthesis pathways in cells by matching precursor supply with utilization.

In summary, we provide evidence that the essential and broadly conserved MraY step in PG synthesis is subject to a previously unknown regulatory mechanism. Mutational and structural evidence identified the likely regulatory site on the enzyme. Importantly, this site is accessible by small molecules from the extracytoplasmic side of the membrane unlike the active site, which is in the cytoplasm. This regulatory site therefore represents an attractive new target for the development of small molecule inhibitors of MraY for potential use as antibiotics.

## ACKNOWLEDGEMENTS

We would like to thank all the members of the Bernhardt, Clemons, Stansfeld, and Rudner Labs for their thoughtful discussions and advice throughout this project. We are also grateful to the members of the CMCB (Center for Microbial Chemical Biology) team at McMaster University and to the Andres and Whitney labs at McMaster University for use of their lab space and facilities by LSM. We thank Lori Burrows for the gift of the B-band serotype O5 monoclonal antibody. This work was supported by the National Institutes of Health (AI083365 to TGB and R01GM114611 to WMC), The G. Harold and Leila Y. Mathers Foundation (to WMC), Wellcome Trust (208361/Z/17/Z to PJS), MRC (MR/S009213/1 to PJS), BBSRC (BB/P01948X/1, BB/R002517/1 and BB/S003339/1 to PJS), the Howard Dalton Centre (to PJS), and Investigator funds from the Howard Hughes Medical Institute (to TGB). LSM was supported by an NSERC Postdoctoral fellowship. DS was supported by a CIHR postdoctoral fellowship. AFG was supported by an NSERC USRA. BWAB’s studentship is sponsored by the MRC. (Cryo)Electron microscopy by WMC’s lab was done in the Beckman Institute Resource Center for Transmission Electron Microscopy at Caltech with help from Songye Chen. MD analysis in PJS’s group made use of time on ARCHER and JADE granted via the UK High-End Computing Consortium for Biomolecular Simulation, HECBioSim (http://hecbiosim.ac.uk), supported by EPSRC (grant no. EP/R029407/1). PJS acknowledges Sulis at HPC Midlands+, which was funded by the EPSRC on grant EP/T022108/1, and the University of Warwick Scientific Computing Research Technology Platform for computational access.

## Data Availability

The atomic coordinates presented in this study have been deposited in the RSCB Protein Data Bank under the accession number PDB: 8TLU.

---

**Supplementary Figure 1.**
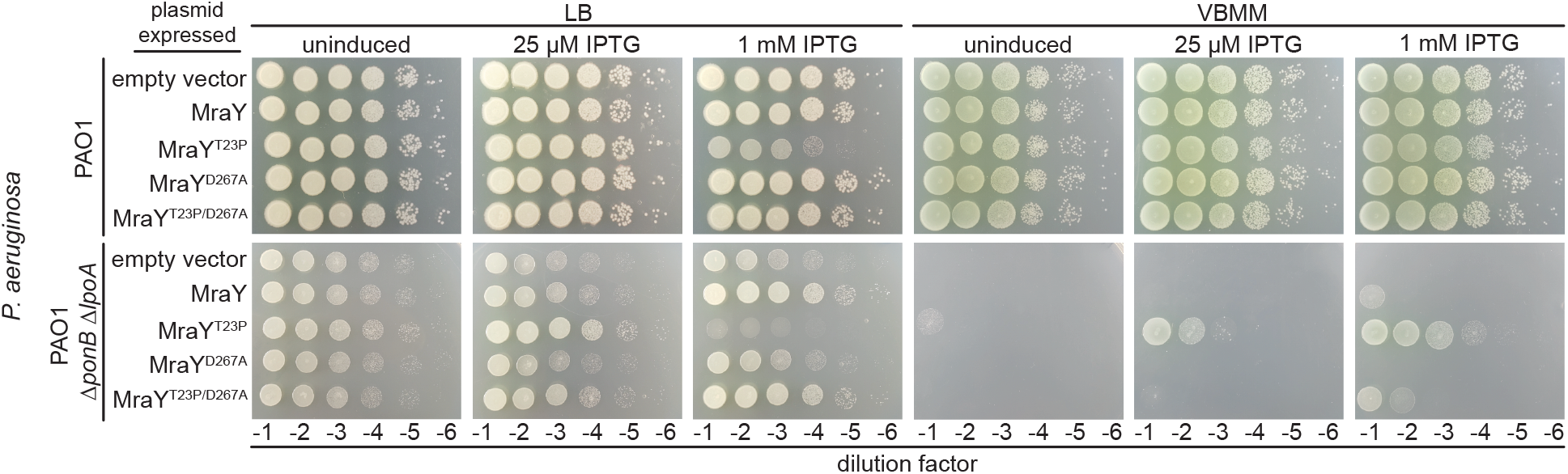
Catalytic activity is required for MraY(T23P) to suppress cell wall defects. Ten-fold serial dilutions of cells of the indicated *P. aeruginosa* strains harboring expression plasmids producing the indicated MraY variant were plated on media with or without IPTG to induce production of MraY variants as indicated.

**Supplementary Figure 2.**
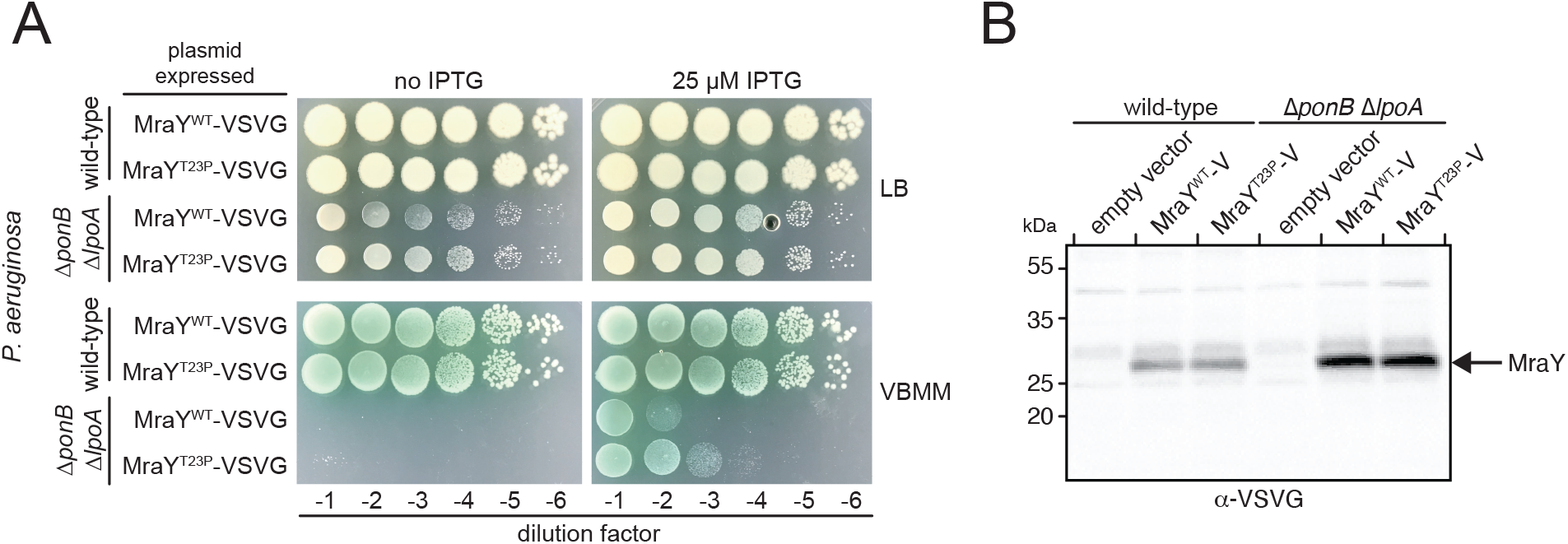
**Cells produce MraY(WT) and MraY(T23P) to comparable levels.** (**A**) Ten-fold serial dilutions of *P. aeruginosa* cells harboring expression plasmids producing the indicated VSVG-tagged MraY were plated on media with or without inducer as indicated. (**B**) Western blot of cells expressing MraY(WT)-VSVG or MraY(T23P)-VSVG. *P. aeruginosa* cells expressing the indicated plasmid were grown to mid-log, normalized for optical density, and extracts were prepared for immunoblotting. Protein was detected using α-VSVG antibody.

**Supplementary Figure 3:**
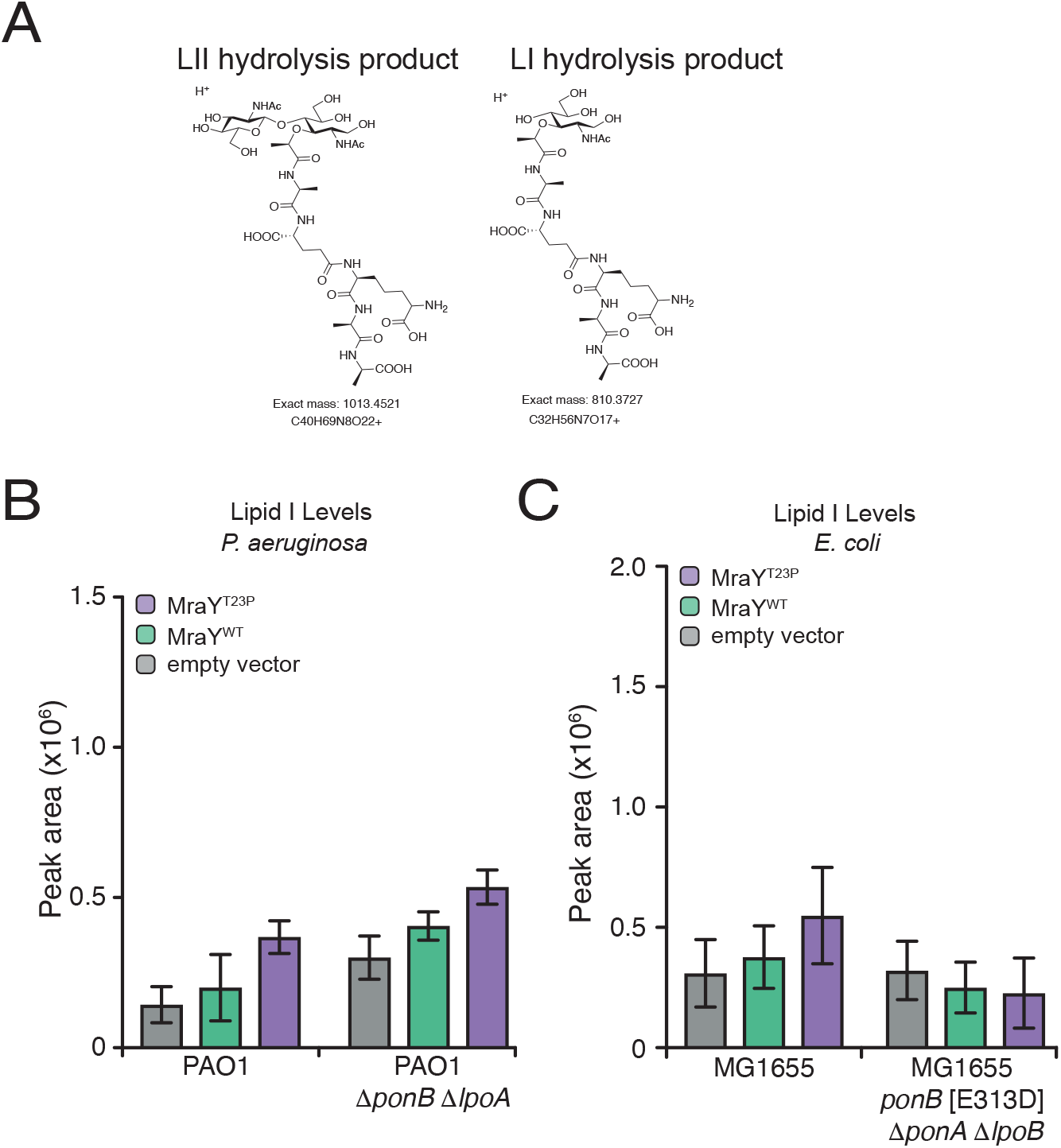
Lipid I levels in cells producing MraY(WT) or MraY(T23P). (**A**) Chemical structures of the Lipid II (LII) and Lipid I (LI) hydrolysis products detected by LCMS. Quantification of extracted ion chromatograms of the lipid I hydrolysis product for the indicated *P. aeruginosa* (**B**) and *E. coli* (**C**) strains. Three independent extractions were performed with lipid I levels quantified using the area of the peak from the extracted ion chromatogram using the Agilent software. Error bars represent SD. For MraY(T23P) vs MraY(WT) in PAO1 1′*ponB* 1′*lpoA* P<0.05, in PAO1, MG1655, MG1655 1′*ponA* 1′*lpoB ponB*[E313D], not significant.

**Supplementary Figure 4:**
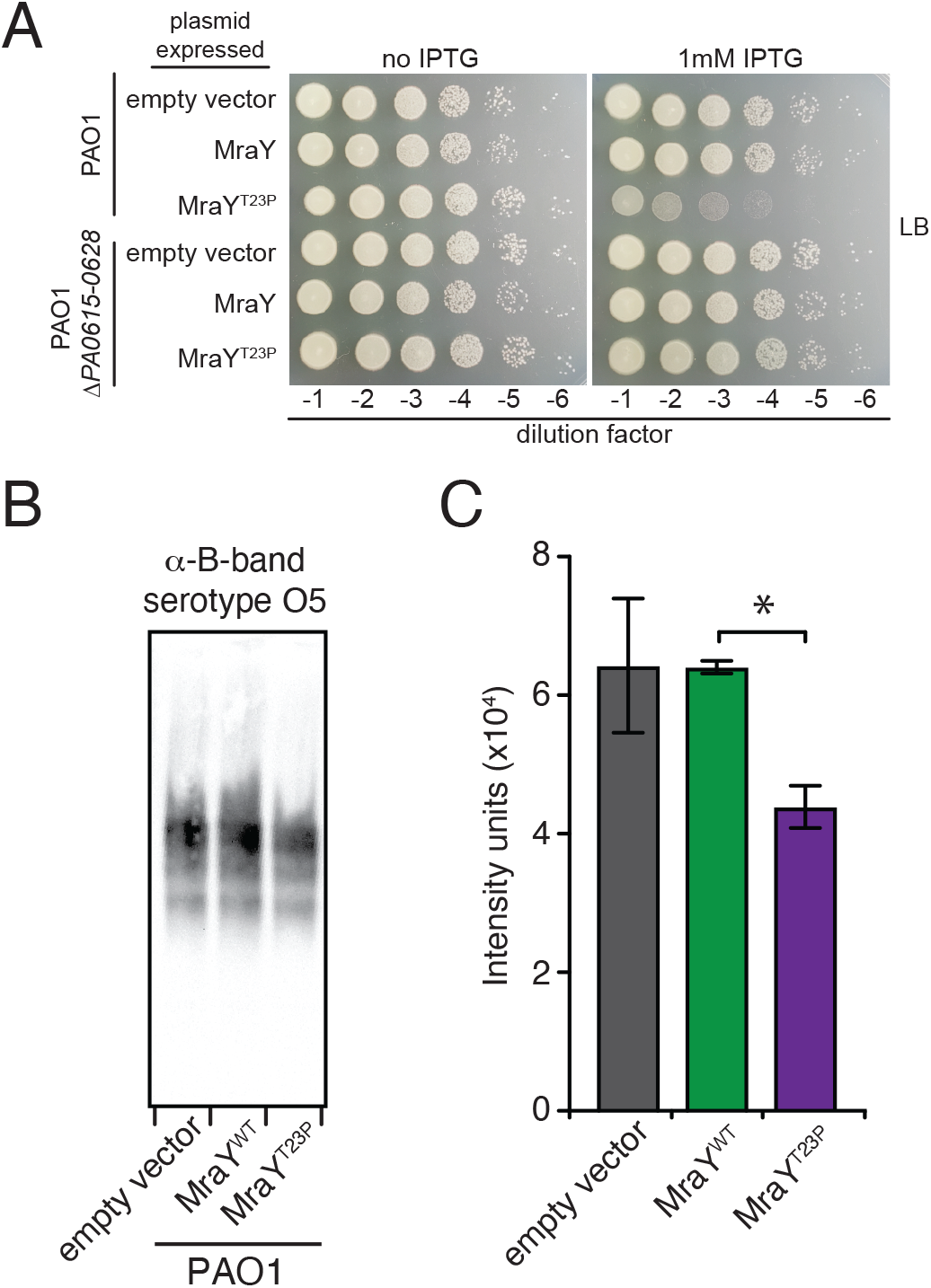
Expression of *^Pa^*MraY(T23P) causes a pyocin-dependent growth defect in *P. aeruginosa* due to a reduction in O-antigen production. (A) Ten-fold serial dilutions of *P. aeruginosa* strains harboring expression plasmids producing the indicated MraY variant were plated on LB containing with or without IPTG to induce protein production from the plasmids. (B) Western blot of B-band O-antigen from *P. aeruginosa* cells expressing the MraY proteins as indicated. Image is representative of three independent experiments. (C) The B-band LPS from three independent replicates of sample extraction was quantified using densitometry. Error bars represent SEM, P <0.05.

**Supplementary Figure 5.**
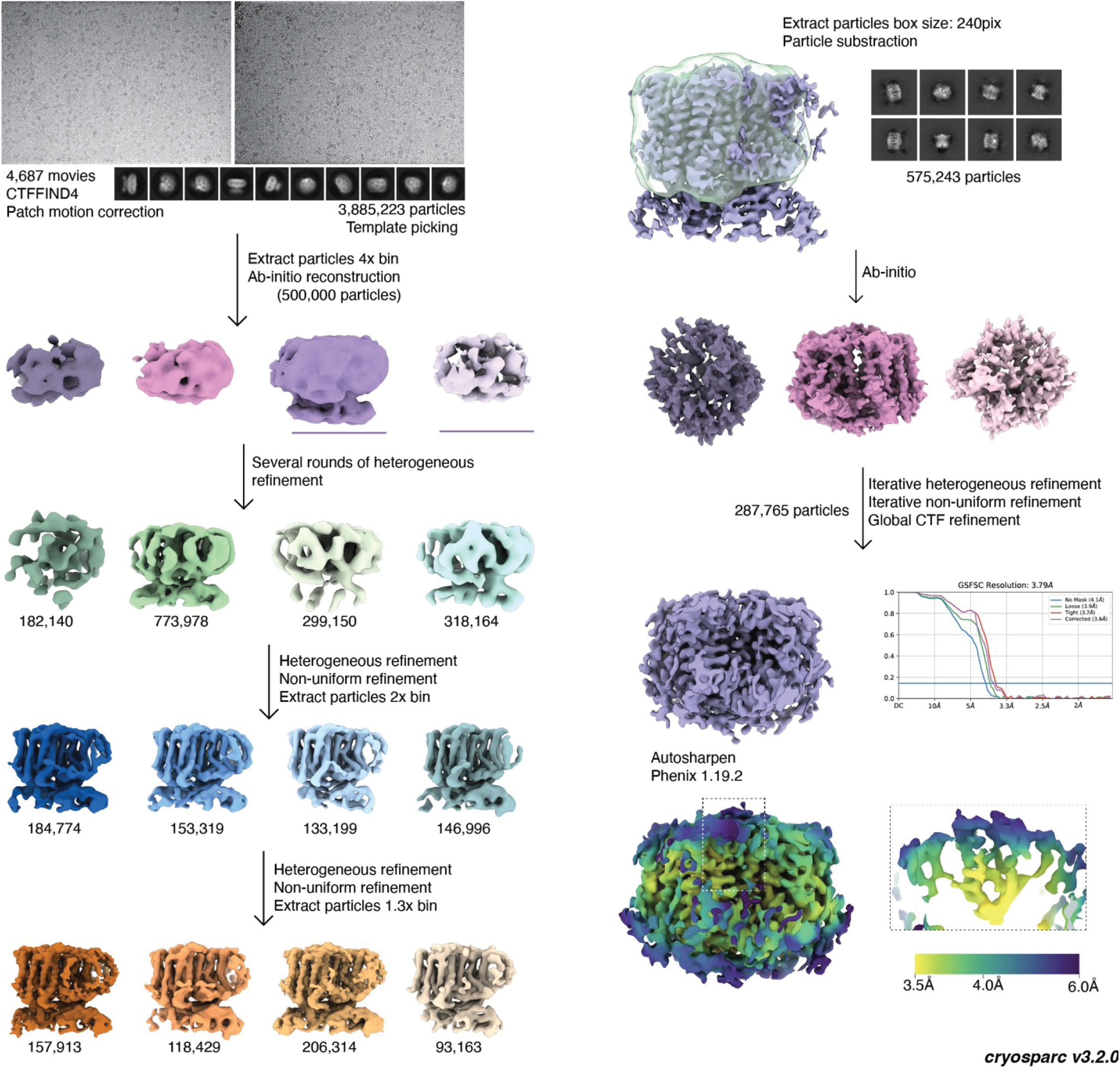
Cryo-EM structure of *^Ec^*MraY(T23P) in the YES complex. Data processing was performed using cryosparc (v3.2.0). Representative movies are shown (top left) with corresponding 2D classes observed in the dataset. Arrows denote the methodology order, following several rounds of heterogeneous refinement. The number of particles sorted is shown below the densities. The masked volume of MraY (green, top right) used for particle subtraction is shown overlayed with the density (purple) of the entire YES complex. The final model is colored by resolution using the viridis color scheme. The unmodeled density at the dimer-interface is isolated for clarity and shown in a dotted box.

**Supplementary Figure 6.**
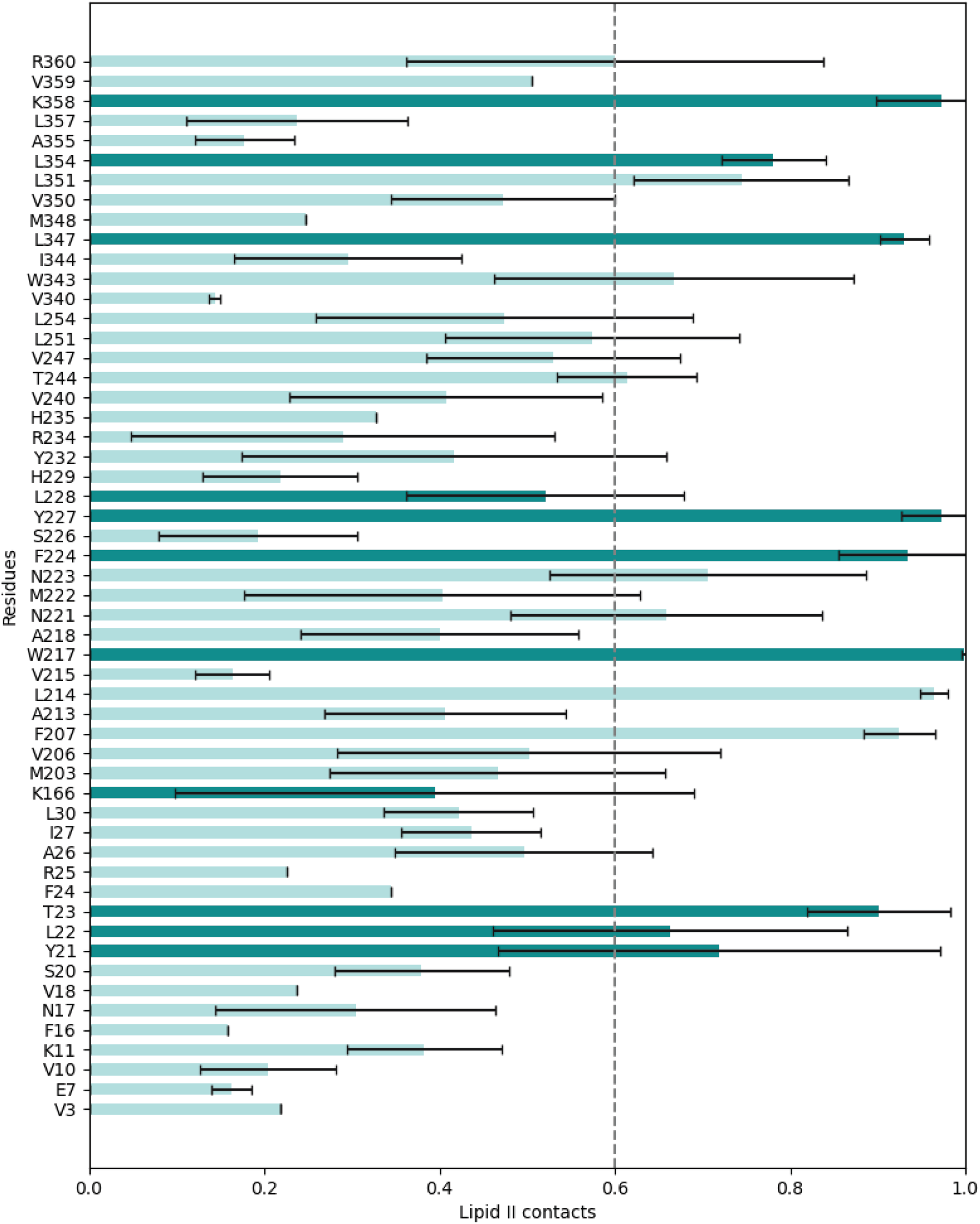
MraY residues contacting lipid II in the MD simulations. Lipid II contacts with MraY residues from atomistic MD simulations. Error bars represent standard error from 5 repeats. Darker green bars represent residues altered in hyperactive variants. Dashed line at x=0.6 represents cutoff for interactions shown in Figure 4C.

**Supplementary Figure 7.**
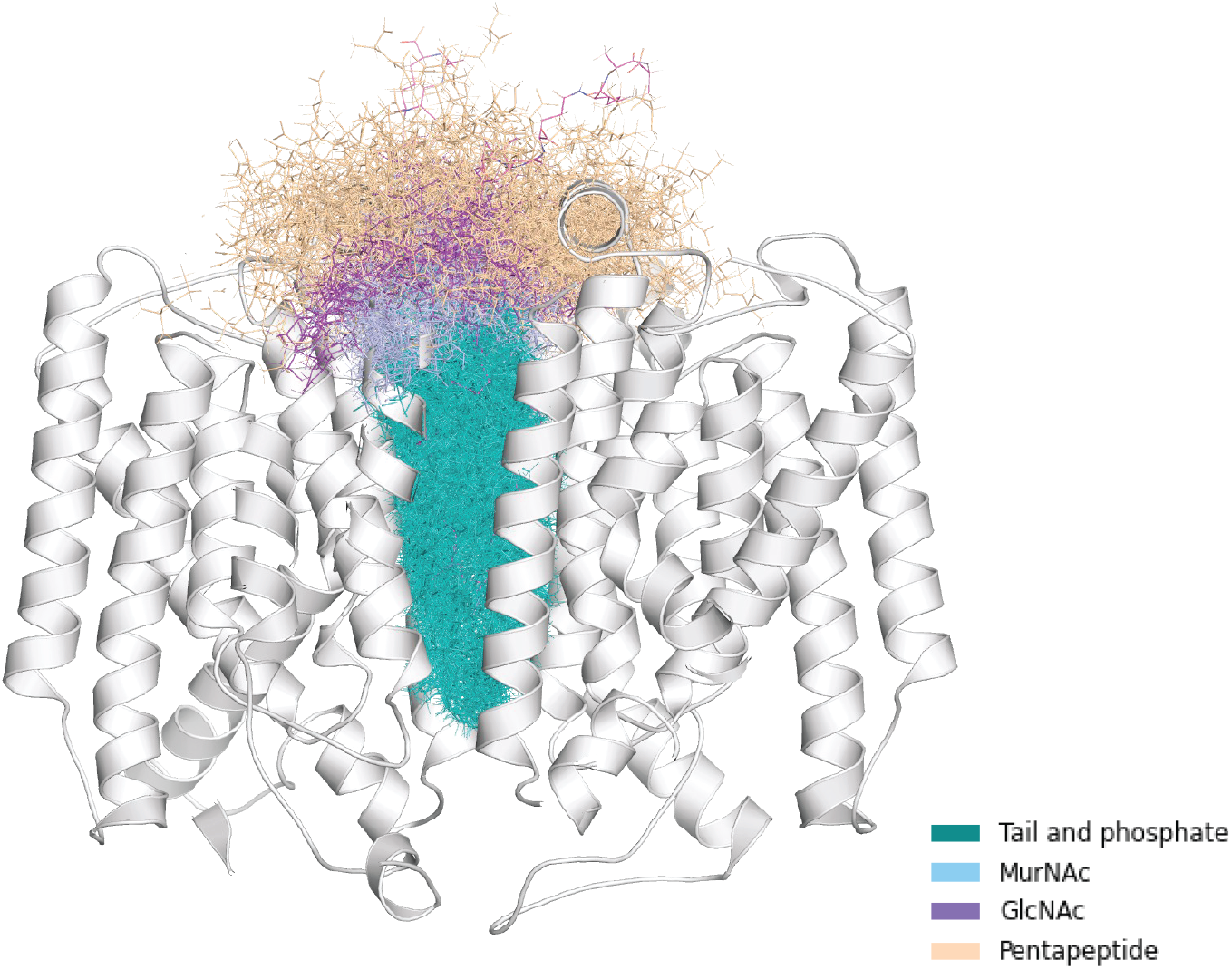
Flexibility of MraY bound lipid II in the MD simulation. All states of lipid II from 5 repeats of atomistic simulation overlaid onto the structure of MraY. Colored as in Fig. 4D.

**Supplementary Figure 8.**
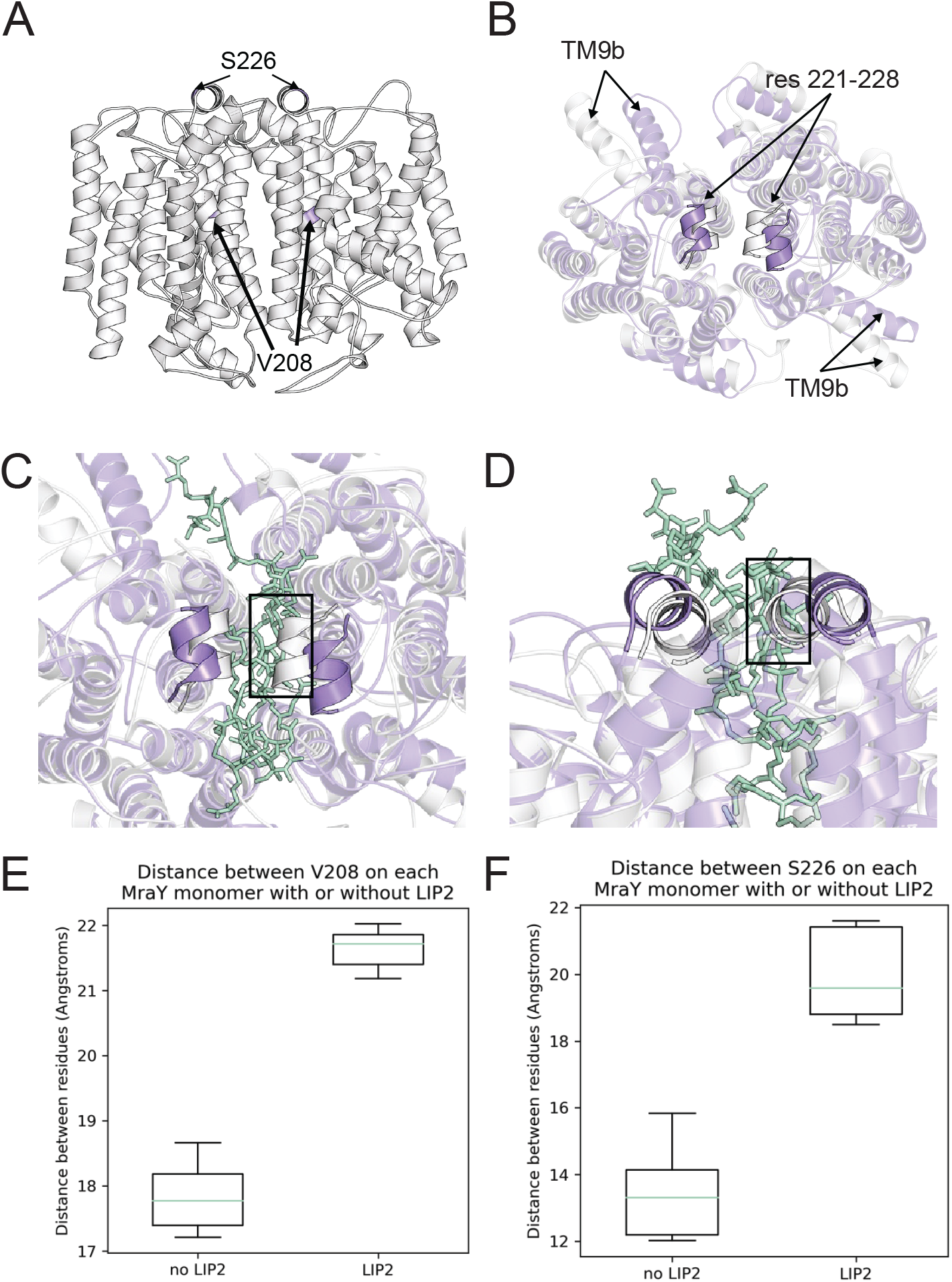
MD analysis identifies potential conformational changes in MraY upon lipid II binding. (A) Structure of MraY dimer in state when lipid II is bound (not shown). Residues V208 and S226 are indicated and colored purple. **(B-D)** An overlay of the structure of MraY at the end of simulations with (purple) or without (gray) lipid II present. **(B)** The structure is shown from the top, lipid II is hidden, and helices with notable differences are indicated. **(C, D)** MraY with lipid II, boxes indicate where lipid II clashes with the structure from the simulation without lipid II, indicating why the periplasmic helix 221-228 is moved apart when lipid II is bound. **(C)** is top (periplasmic) view, while **(D)** is a side view. **(E)** A boxplot of the average distance between V208 (a residue in the lipid II binding pocket) of each MraY monomer, in simulations with or without lipid II present. The data represented by each box plot is the mean distance from all time points in each of 5 repeats. **(F)** A boxplot of the average distance between S226 (a residue in the periplasmic helix above the lipid II binding site) of each MraY monomer, in simulations with or without lipid II present. The data represented by each box plot is the mean distance from all time points in each of 5 repeats.

**Supplementary Figure 9.**
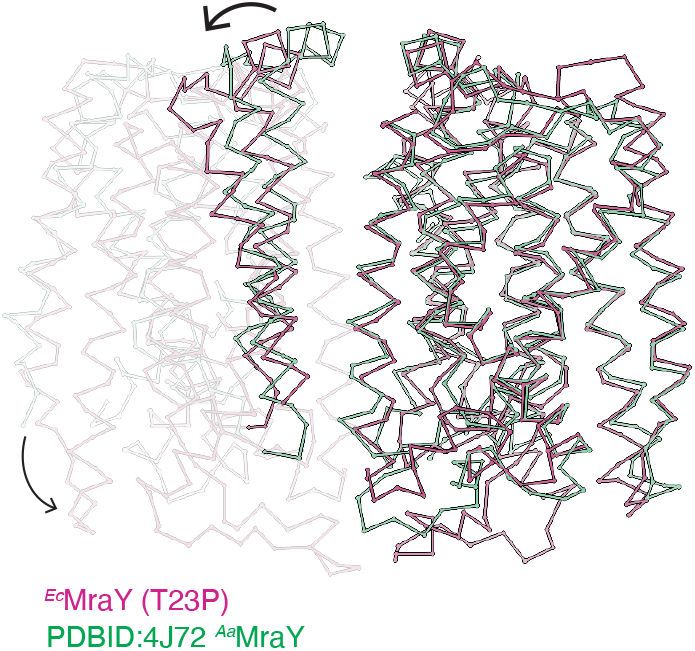
Altered conformation of MraY dimers in the YES complex versus *^Aa^*MraY. View from the plane of the membrane. Stick representation of the α-carbon chain of *^Ec^*MraY(T23P) (pink) structurally aligned to *^Aa^*MraY (PDBID:4J72)(green). Molecules are aligned to the right chains in the figure. Arrows highlight the differences in *^Aa^*MraY compared to *^Ec^*MraY(T23P).

**Supplementary Figure 10.**
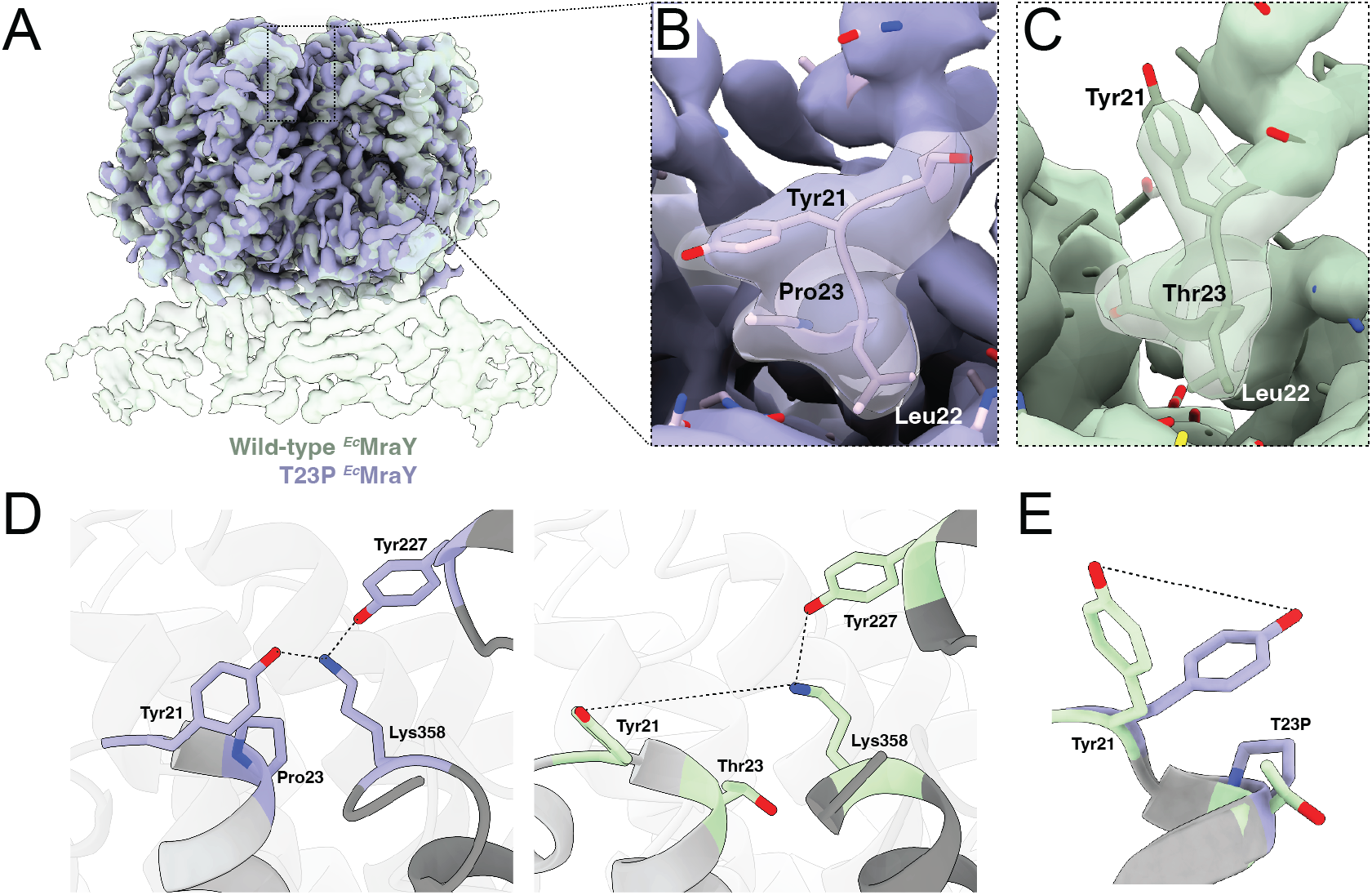
Comparison of MraY(WT) and MraY(T23P) structures in the YES complex. (**A**) Overlay of densities of MraY(WT) (EMDB-29641) (green) and MraY(T23P) (purple) viewed in the plane of the membrane. (**B**) Enlarged view of the densities around the T23P mutant. Residues are shown in stick representation. Residues 21-23 are labeled for reference. (**C**) As in *B* for the wild-type complex. (**D**) Hydrogen bonding network observed in MraY(T23P) (left, purple) compared to WT (right, green) at the mutagenesis site involving Y21, Y227, and K358. (**E**) Similar to *D,* overlay of the two models highlighting the conformational differences of residue Y21.

**Supplementary Movie 1 : Movie of Lipid II binding MraY dimer**. 10 µs simulation of the top view of the MraY dimer in a mixed CG membrane. Representative simulation of two lipid II molecules (highlighted as green, gold, and pink spheres) freely entering the MraY cavity during unbiased MD simulations.

**Supplementary Movie 2 : Movie of C55P binding MraY dimer.** 10 µs simulation of the top view of the MraY dimer in a mixed CG membrane. One C55P molecule (highlighted as green and gold spheres) freely enters the MraY cavity during unbiased MD simulations when lipid I and lipid II are omitted.

## Supplementary Material

**SI Table 1.** List of suppressing MraY variants.

| Residue | Changed to |
| --- | --- |
| K14 | E |
| Y21 | H |
| L22 | P |
| T23 | A, P |
| L31 | P |
| L160 | R |
| M164 | R |
| K166 | E |
| Y217 | H |
| G220 | S |
| F224 | S |
| A225 | V |
| Y227 | C |
| L228 | P |
| G248 | C |
| Q286 | R |
| V289 | M |
| I339 | V |
| V347 | M |
| L354 | P |
| K358 | M |

**SI Table 2:** Cryo-EM data collection, refinement, and validation statistics *Ec*MraY.

| ***EcMraY*** |  |
| --- | --- |
| (EMDB- EMD-41373) |  |
| (PDB 8TLU) |  |
| **Data collection and processing** |  |
| Microscope | FEI Titan Krios |
| Magnification | 105,000 |
| Voltage (kV) | 300 |
| Electron exposure (e-/Å<sup>2</sup>) | 60 |
| Defocus range (µm) | -1 to -2.5 |
| Pixel size (Å) | 0.832 |
| Symmetry imposed | C1 |
| Initial particle images (no.) | 3,885,223 |
| Final particle images (no.) | 287,765 |
| Map resolution (Å) | 3.8 |
| FSC threshold: 0.143 |  |
| **Refinement** |  |
| Software | PHENIX 1.19.2 |
| Initial model used (PDB code) | 8G01 |
| Resolution of unmasked reconstructions (Å, FSC=0.5) | 4.1 |
| Resolution of masked reconstructions (Å, FSC=0.5) | 3.8 |
| Correlation coefficient ($CC_{mask}$) | 0.76 |
| Model composition |  |
| Atoms (Hydrogens) | 11180 (5703) |
| Protein residues | 698 |
| Ligands | 0 |
| *B* factors (Å<sup>2</sup>) (min/max/mean) |  |
| Protein | 20.34/79.77/41.69 |
| Ligand | - |
| R.m.s. deviations |  |
| Bond lengths (Å) | 0.003 (0) |
| Bond angles (°) | 0.536 (0) |
| **Validation** |  |
| MolProbity score | 1.43 |
| Clashscore | 7.87 |
| Poor rotamers (%) | 0 |
| Ramachandran plot |  |
| Favored (%) | 98.40 |
| Allowed (%) | 1.60 |
| Disallowed (%) | 0 |

**SI Table 3.**
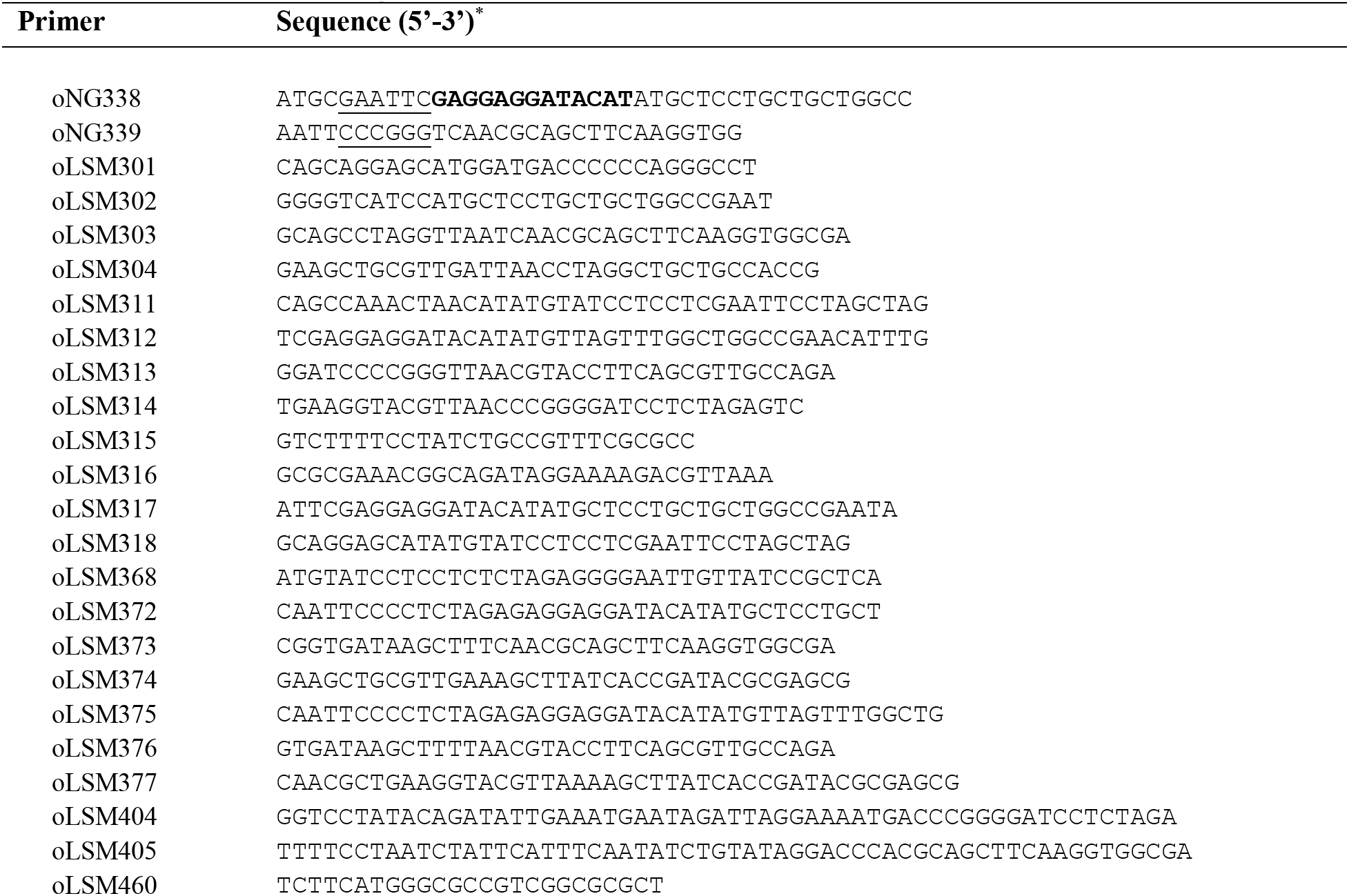
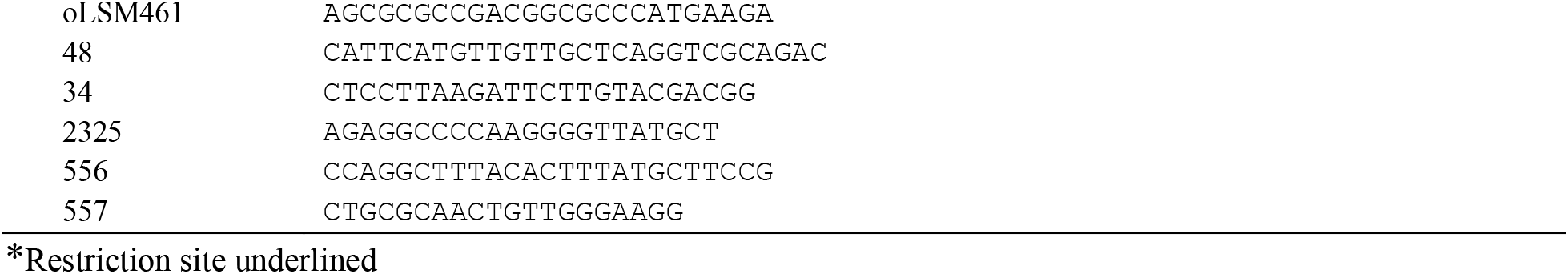
Oligonucleotide primers used in this study.

**SI Table 4.** Bacterial strains used in this study.

| Strain | Description* | Reference |
| --- | --- | --- |
| *E. coli* |  |  |
| DH5α | Host strain for plasmid cloning | Invitrogen |
| BL21(DE3) | Expression strain for protein production | 1 |
| CAM333 | C43(DE3) ΔponB, ΔpbpC, ΔmtgA | 2 |
| LSM9 | C43(DE3) ΔpbpC, ΔmtgA *fhuA*- | This study |
| ΔslyD BL21(DE3) | | 3 |
| MG1655 | Wild-type | 4 |
| MM119 | ΔponA *ponB*(E313D) ΔlpoB | 5 |
| *P. aeruginosa* |  |  |
| PAO1 | Wild-type | S. Lory |
| PA686 | ΔponB ΔlpoA | 6 |
| PA760 | ΔponB ΔlpoA *mraY*(T23P) | This study |
| PA662 | ΔPA0615–0628, R2 pyocin deletion | S. Lory |

**SI Table 5.**
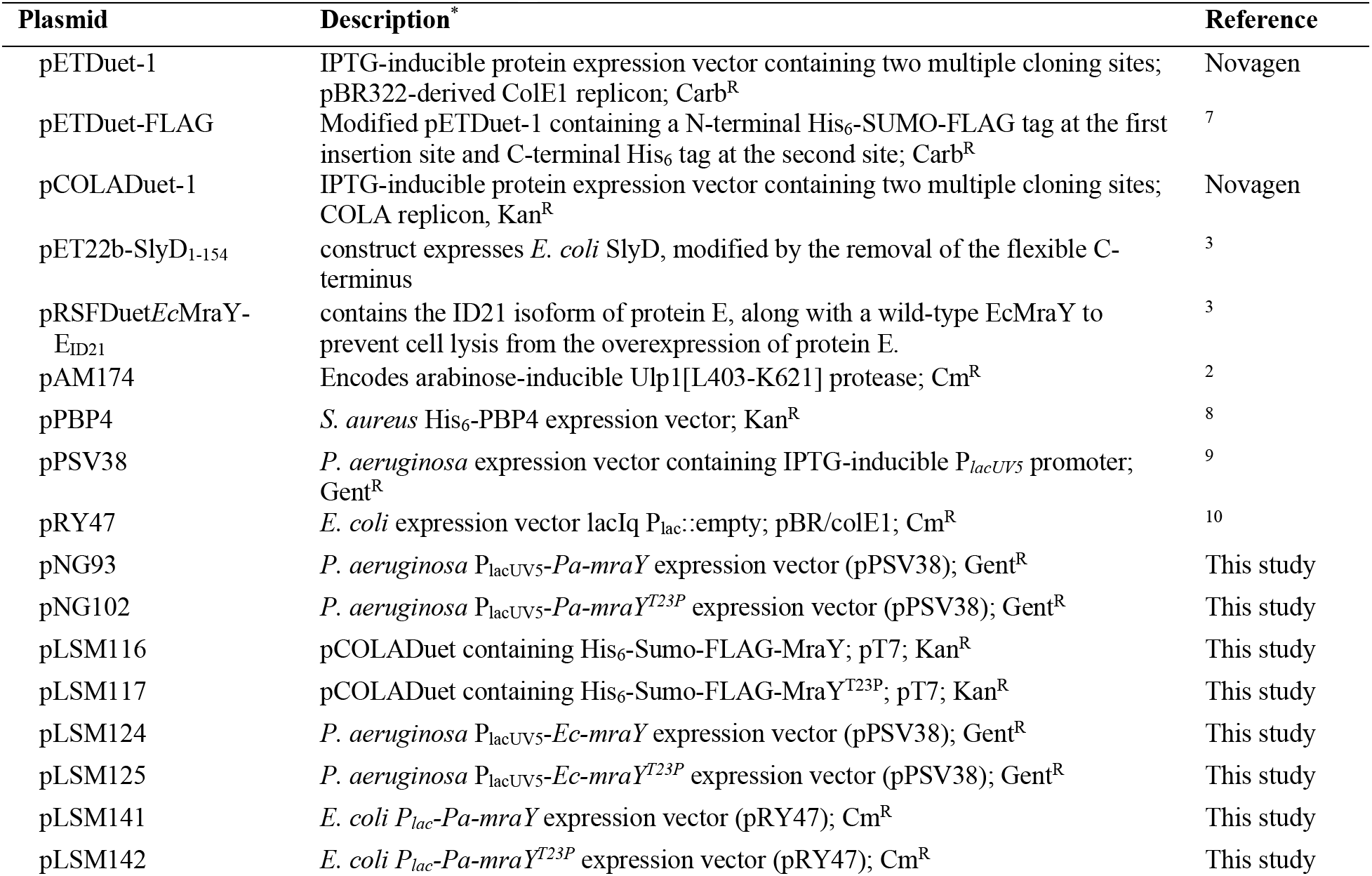
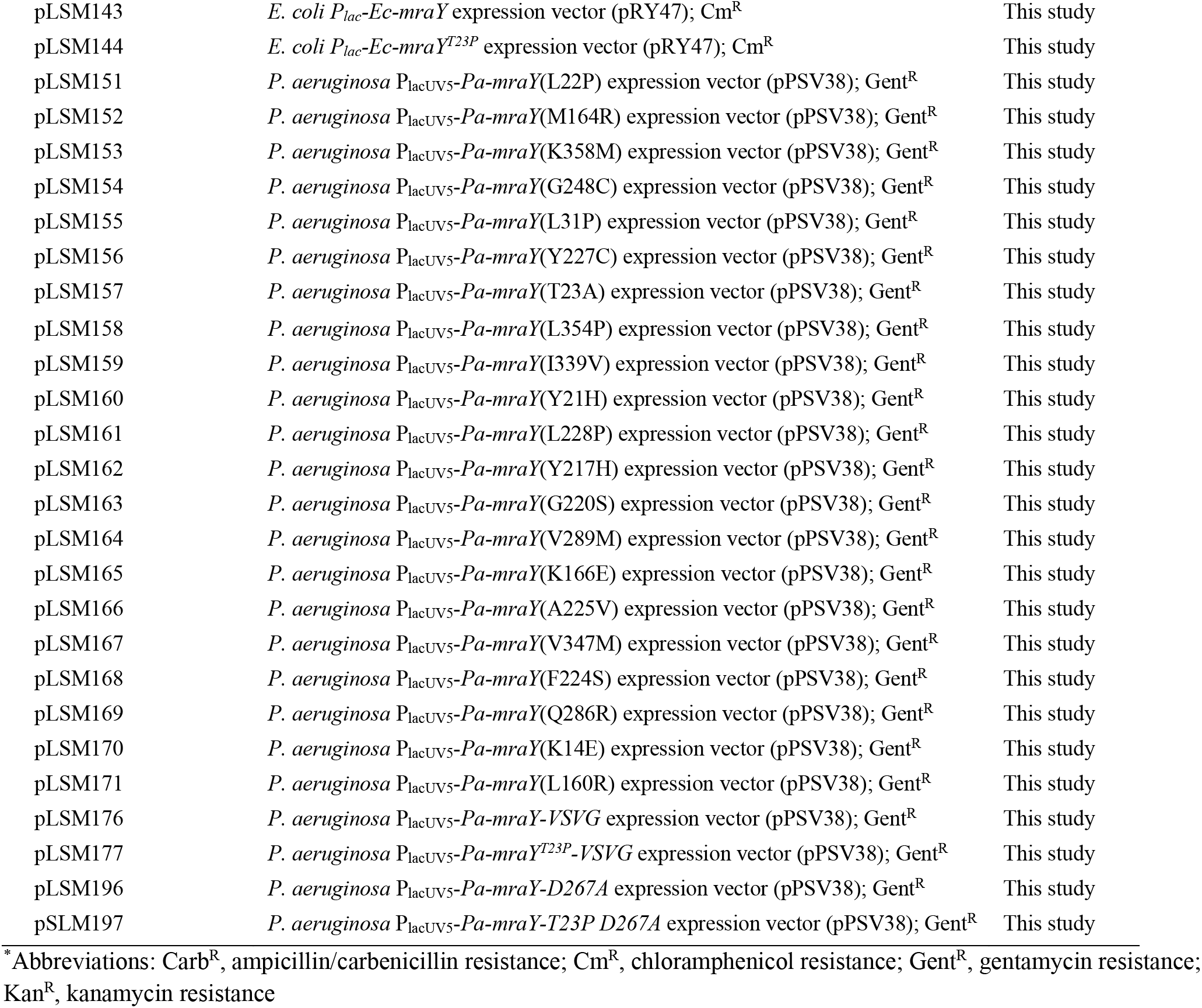
Plasmids used in this study **Plasmid Description**\*.

| Plasmid | Description* | Reference |
| --- | --- | --- |
| pETDuet-1 | IPTG-inducible protein expression vector containing two multiple cloning sites; pBR322-derived ColE1 replicon; Carb<sup>R</sup> | Novagen |
| pETDuet-FLAG | Modified pETDuet-1 containing a N-terminal His<sub>6</sub>-SUMO-FLAG tag at the first insertion site and C-terminal His<sub>6</sub> tag at the second site; Carb<sup>R</sup> | 7 |
| pCOLADuet-1 | IPTG-inducible protein expression vector containing two multiple cloning sites; COLA replicon, Kan<sup>R</sup> | Novagen |
| pET22b-SlyD<sub>1-154</sub> | construct expresses *E. coli* SlyD, modified by the removal of the flexible C-terminus | 3 |
| pRSFDuetEcMraY-E<sub>ID21</sub> | contains the ID21 isoform of protein E, along with a wild-type EcMraY to prevent cell lysis from the overexpression of protein E. | 3 |
| pAM174 | Encodes arabinose-inducible Ulp1[L403-K621] protease; Cm<sup>R</sup> | 2 |
| pPBP4 | *S. aureus* His<sub>6</sub>-PBP4 expression vector; Kan<sup>R</sup> | 8 |
| pPSV38 | *P. aeruginosa* expression vector containing IPTG-inducible P<sub>lacUV5</sub> promoter; Gent<sup>R</sup> | 9 |
| pRY47 | *E. coli* expression vector lacIq P<sub>lac</sub>::empty; pBR/colE1; Cm<sup>R</sup> | 10 |
| pNG93 | *P. aeruginosa* P<sub>lacUV5</sub>-*Pa-mraY* expression vector (pPSV38); Gent<sup>R</sup> | This study |
| pNG102 | *P. aeruginosa* P<sub>lacUV5</sub>-*Pa-mraY*<sup>T23P</sup> expression vector (pPSV38); Gent<sup>R</sup> | This study |
| pLSM116 | pCOLADuet containing His<sub>6</sub>-Sumo-FLAG-MraY; pT7; Kan<sup>R</sup> | This study |
| pLSM117 | pCOLADuet containing His<sub>6</sub>-Sumo-FLAG-MraY<sup>T23P</sup>; pT7; Kan<sup>R</sup> | This study |
| pLSM124 | *P. aeruginosa* P<sub>lacUV5</sub>-*Ec-mraY* expression vector (pPSV38); Gent<sup>R</sup> | This study |
| pLSM125 | *P. aeruginosa* P<sub>lacUV5</sub>-*Ec-mraY*<sup>T23P</sup> expression vector (pPSV38); Gent<sup>R</sup> | This study |
| pLSM141 | *E. coli* P<sub>lac</sub>-*Pa-mraY* expression vector (pRY47); Cm<sup>R</sup> | This study |
| pLSM142 | *E. coli* P<sub>lac</sub>-*Pa-mraY*<sup>T23P</sup> expression vector (pRY47); Cm<sup>R</sup> | This study |
| pLSM143 | *E. coli* P<sub>lac</sub>-*Ec-mraY* expression vector (pRY47); Cm<sup>R</sup> | This study |
| pLSM144 | *E. coli* P<sub>lac</sub>-*Ec-mraY*<sup>T23P</sup> expression vector (pRY47); Cm<sup>R</sup> | This study |
| pLSM151 | *P. aeruginosa* P<sub>lacUV5</sub>-*Pa-mraY*(L22P) expression vector (pPSV38); Gent<sup>R</sup> | This study |
| pLSM152 | *P. aeruginosa* P<sub>lacUV5</sub>-*Pa-mraY*(M164R) expression vector (pPSV38); Gent<sup>R</sup> | This study |
| pLSM153 | *P. aeruginosa* P<sub>lacUV5</sub>-*Pa-mraY*(K358M) expression vector (pPSV38); Gent<sup>R</sup> | This study |
| pLSM154 | *P. aeruginosa* P<sub>lacUV5</sub>-*Pa-mraY*(G248C) expression vector (pPSV38); Gent<sup>R</sup> | This study |
| pLSM155 | *P. aeruginosa* P<sub>lacUV5</sub>-*Pa-mraY*(L31P) expression vector (pPSV38); Gent<sup>R</sup> | This study |
| pLSM156 | *P. aeruginosa* P<sub>lacUV5</sub>-*Pa-mraY*(Y227C) expression vector (pPSV38); Gent<sup>R</sup> | This study |
| pLSM157 | *P. aeruginosa* P<sub>lacUV5</sub>-*Pa-mraY*(T23A) expression vector (pPSV38); Gent<sup>R</sup> | This study |
| pLSM158 | *P. aeruginosa* P<sub>lacUV5</sub>-*Pa-mraY*(L354P) expression vector (pPSV38); Gent<sup>R</sup> | This study |
| pLSM159 | *P. aeruginosa* P<sub>lacUV5</sub>-*Pa-mraY*(I339V) expression vector (pPSV38); Gent<sup>R</sup> | This study |
| pLSM160 | *P. aeruginosa* P<sub>lacUV5</sub>-*Pa-mraY*(Y21H) expression vector (pPSV38); Gent<sup>R</sup> | This study |
| pLSM161 | *P. aeruginosa* P<sub>lacUV5</sub>-*Pa-mraY*(L228P) expression vector (pPSV38); Gent<sup>R</sup> | This study |
| pLSM162 | *P. aeruginosa* P<sub>lacUV5</sub>-*Pa-mraY*(Y217H) expression vector (pPSV38); Gent<sup>R</sup> | This study |
| pLSM163 | *P. aeruginosa* P<sub>lacUV5</sub>-*Pa-mraY*(G220S) expression vector (pPSV38); Gent<sup>R</sup> | This study |
| pLSM164 | *P. aeruginosa* P<sub>lacUV5</sub>-*Pa-mraY*(V289M) expression vector (pPSV38); Gent<sup>R</sup> | This study |
| pLSM165 | *P. aeruginosa* P<sub>lacUV5</sub>-*Pa-mraY*(K166E) expression vector (pPSV38); Gent<sup>R</sup> | This study |
| pLSM166 | *P. aeruginosa* P<sub>lacUV5</sub>-*Pa-mraY*(A225V) expression vector (pPSV38); Gent<sup>R</sup> | This study |
| pLSM167 | *P. aeruginosa* P<sub>lacUV5</sub>-*Pa-mraY*(V347M) expression vector (pPSV38); Gent<sup>R</sup> | This study |
| pLSM168 | *P. aeruginosa* P<sub>lacUV5</sub>-*Pa-mraY*(F224S) expression vector (pPSV38); Gent<sup>R</sup> | This study |
| pLSM169 | *P. aeruginosa* P<sub>lacUV5</sub>-*Pa-mraY*(Q286R) expression vector (pPSV38); Gent<sup>R</sup> | This study |
| pLSM170 | *P. aeruginosa* P<sub>lacUV5</sub>-*Pa-mraY*(K14E) expression vector (pPSV38); Gent<sup>R</sup> | This study |
| pLSM171 | *P. aeruginosa* P<sub>lacUV5</sub>-*Pa-mraY*(L160R) expression vector (pPSV38); Gent<sup>R</sup> | This study |
| pLSM176 | *P. aeruginosa* P<sub>lacUV5</sub>-*Pa-mraY*-*VSVG* expression vector (pPSV38); Gent<sup>R</sup> | This study |
| pLSM177 | *P. aeruginosa* P<sub>lacUV5</sub>-*Pa-mraY*<sup>T23P</sup>-*VSVG* expression vector (pPSV38); Gent<sup>R</sup> | This study |
| pLSM196 | *P. aeruginosa* P<sub>lacUV5</sub>-*Pa-mraY*-*D267A* expression vector (pPSV38); Gent<sup>R</sup> | This study |
| pSLM197 | *P. aeruginosa* P<sub>lacUV5</sub>-*Pa-mraY*-*T23P D267A* expression vector (pPSV38); Gent<sup>R</sup> | This study |

\*Abbreviations: Carb<sup>R</sup>, ampicillin/carbenicillin resistance; Cm<sup>R</sup>, chloramphenicol resistance; Gent<sup>R</sup>, gentamycin resistance; Kan<sup>R</sup>, kanamycin resistance

## SI METHODS

### Plasmid construction

#### pNG93 [P_lacUV5_::*Pa-mraY*(*PA4415*)] is a pPSV38 derivative

pPSV38 was digested with EcoRI/XmaI to generate the plasmid backbone. *P. aeruginosa mraY* (PA4415; M1-R360) was amplified from PAO1 gDNA with oNG338/oNG339 to introduce a synthetic RBS (5’ - GAGGAGGATACAT - 3’). After digestion with EcoRI/XmaI, the PCR product was ligated into pPSV38 to generate pNG93. The final construct was sequence verified using primers 556 and 557. The *mraY* gene in this and all related constructs below is inducible with IPTG.

#### pNG102 [P_lacUV5_::*Pa-mraY(T23P)*] is a pPSV38 derivative

pPSV38 was digested with EcoRI/XmaI to generate the plasmid backbone. *P. aeruginosa mraY* T23P was amplified from PA760 via colony PCR with oNG338/oNG339 to introduce a synthetic RBS (5’ - GAGGAGGATACAT - 3’).). After digestion with EcoRI/XmaI, the PCR product was ligated into pPSV38 to generate pNG93. The final construct was sequence verified using primers 556 and 557.

#### pLSM116 [P_T7_::*H-SUMO-FLAG-Pa-mraY*] is a pCOLADuet derivative

The gene encoding full-length *P. aeruginosa mraY* was amplified from pNG93 using the primers oLSM302 and oLSM303. Using pCOLADuet as a template, the backbone was amplified using oLSM301 and oLSM304. The fragments were joined using Gibson assembly and sequence verified using primers 34 and 2325.

#### pLSM117 [P_T7_::*H-SUMO-FLAG-Pa-mraY^T23P^*] is a pCOLADuet derivative

The gene encoding full-length *P. aeruginosa mraY^T23P^* was amplified from pNG102 using the primers oLSM302 and oLSM303. Using pCOLADuet as a template, the backbone was amplified using oLSM301 and oLSM304. The fragments were joined using Gibson assembly and sequence verified using primers 34 and 2325.

#### pLSM124 [P_lacUV5_::*Ec-mraY*] is a pPSV38 derivative

The gene encoding full-length *E. coli mraY* was amplified from MG1655 genomic DNA using primers oLSM312 and oLSM313. Using pNG93 as a template, the backbone was amplified using oLSM311 and oLSM314. The fragments were joined using Gibson assembly. The final construct was sequence verified using primers 556 and 557.

#### pLSM125 [P_lacUV5_::*Ec-mraY(T23P)*] is a pPSV38 derivative

Using pLSM124 as a template, T23 was mutated to P using site directed mutagenesis (QuikChange Lightning, Agilent) using the primers oLSM315 and oLSM316. The final construct was sequence verified using primers 556 and 557.

#### pLSM141 [P_lac_::*Pa*-*mraY*] is a pRY47 derivative

The gene encoding full-length *P. aeruginosa mraY* was amplified from pNG93 using primers oLSM372 and oLSM373. Using pRY47 as a template, the backbone was amplified using oLSM374 and oLSM368. The fragments were joined using Gibson assembly. The final construct was sequence verified using primers 556 and 48.

#### pLSM142 [P_lac_::*Pa*-*mraY(T23P)*] is a pRY47 derivative

The gene encoding full-length *P. aeruginosa mraY* (T23P) was amplified from pNG102 using primers oLSM372 and oLSM373. Using pRY47 as a template, the backbone was amplified using oLSM374 and oLSM368. The fragments were ligated using Gibson assembly. The final construct was sequence verified using primers 556 and 48.

#### pLSM143 [P_lac_::*Ec-mraY*] is a pRY47 derivative

The gene encoding full-length *E. coli mraY* was amplified from pLSM124 using primers oLSM375 and oLSM376. Using pRY47 as a template, the backbone was amplified using oLSM377 and oLSM368. The fragments were ligated using Gibson assembly. The final construct was sequence verified using primers 556 and 48.

#### pLSM144 [P_lac_::*Ec*-*mraY(T23P)*] is a pRY47 derivative

The gene encoding full-length *E. coli mraY* (T23P) was amplified from pLSM125 using primers oLSM375 and oLSM376. Using pRY47 as a template, the backbone was amplified using oLSM377 and oLSM368. The fragments were ligated using Gibson assembly. The final construct was sequence verified using primers 556 and 48.

#### pLSM176 [P_lacUV5_::*Pa-mraY*-GS-VSVG] is a pPSV38 derivative

The gene encoding full-length *P. aeruginosa mraY* was amplified from pNG93 using primers oLSM317 and oLSM405. Using pNG93 as a template, the backbone was amplified using oLSM404 and oLSM318. The fragments were ligated using Gibson assembly. The final construct was sequence verified using primers 556 and 557.

#### pLSM177 [P_lacUV5_::*Pa-mraY(T23P)*-GS-VSVG] is a pPSV38 derivative

The gene encoding full-length *P. aeruginosa mraY* (T23P) was amplified from pNG102 using primers oLSM317 and oLSM405. Using pNG102 as a template, the backbone was amplified using oLSM404 and oLSM318. The fragments were ligated using Gibson assembly. The final construct was sequence verified using primers 556 and 557.

#### pLSM196 [P_lacUV5_::*Pa-mraY*(*D267A)*] is a pPSV38 derivative

Site directed mutagenesis (QuikChange Lightning, Agilent) of pNG93 was performed to make the D267A change using oLSM460 and oLSM461.

#### pLSM197 [P_lacUV5_::*Pa-mraY(T23P/D267A)*] is a pPSV38 derivative

Site directed mutagenesis (QuikChange Lightning, Agilent) of pNG102 was performed to make the D267A change using oLSM460 and oLSM461.

## Materials

Unless otherwise indicated, all chemicals and reagents were purchased from Sigma-Aldrich. Restriction enzymes were purchased from New England Biolabs. Oligonucleotide primers were purchased from Integrated DNA Technologies.

### Bacterial strains, plasmids, oligonucleotide primers and culture conditions

*E. coli* strains were grown with shaking at 37 °C in lysogeny broth (LB, 10 g/L tryptone, 5 g/L NaCl, 5 g/L yeast extract), lysogeny broth with no salt (LBNS; 10 g/L tryptone, 5 g/L yeast extract), Terrific Broth (TB; 12 g/L tryptone, 24 g/L yeast extract, 0.4% v/v glycerol, 0.17 M KH_2_PO_4_, 0.72 M K_2_HPO_4_), or on LB or LBNS agar as indicated. MM119 was grown at 30°C. *P. aeruginosa* strains are all derivatives of PAO1 and were grown with shaking at 37°C in LB, LBNS, Vogel-Bonner minimal media (VBMM; 3.42 g/L trisodium citrate dihydrate, 2.0 g/L citric acid, 10 g/L K_2_HPO_4_, 3.5 g/L NaNH_4_PO_4_•4H_2_O, 1 mM MgSO_4_, 0.1 mM CaCl_2_) or on LB, LBNS, or VBMM agar as indicated. The following concentration of antibiotics were used to maintain plasmids: ampicillin (Amp), 50 µg/mL; chloramphenicol (Cam), 25 µg/mL; gentamicin (Gent), 15 µg/mL (*E. coli*); Gent, 30 µg/mL (*P. aeruginosa*). The primers, strains and plasmids used in this study are summarized in **SI appendix, Tables 3-5**.

### Electroporation of P. aeruginosa

*P. aeruginosa* strains were made competent using previously described methods^11^. For electroporation, 100 ng of plasmid DNA was added to 40 μL of competent *P. aeruginosa* cells. Transformation was achieved using standard protocols and transformants were selected for using 30 μg/mL Gent.

### Viability assays

Overnight cultures of PAO1, PA686, or PA760 derivatives, containing vectors producing the indicated alleles of *mraY* expressed from an IPTG-inducible (P_lacUV5_) plasmid were normalized to an OD_600_ of 2.4 before being serially diluted. Aliquots (5 µL) of the dilutions were spotted onto LB Gent agar, VBMM Gent agar, with or without IPTG. Plates were incubated at 30°C for 24 h at which point the plates were imaged. A similar protocol was adapted for MG1655 and MM119 derivatives containing vectors producing the indicated alleles of *mraY* from an IPTG inducible (P_lac_) plasmid.

### Immunoblotting

For analysis of protein levels from strains producing MraY-VSVG variants, an overnight culture of each of the strains was allowed to grow in LB containing 30 μg/mL Gent at 37°C. The following day, the cultures were diluted to an OD_600_ of 0.01 and allowed to grow at 37°C in LB containing 30 μg/ml Gent. After 2 h, 1 mM IPTG was added and the cultures were allowed to grow for another 2.5 h. Cultures were normalized to an OD_600_ = 1.0 and cells were collected by centrifugation at 5,000 × g for 2 min. The cell pellet was resuspended in 200 μL of 2× Laemmli buffer, then centrifuged for 10 min at 21,000 x g. Samples were analyzed by SDS-PAGE followed by imunoblotting. Protein was transferred from the SDS-PAGE gel to a nitrocellulose membrane using wet transfer (30 min at 100V) in cold transfer buffer (192 mM glycine, 20% methanol, 25 mM Tris base). The membrane was blocked in 5% (w/v) skim milk powder in Tris-Buffered saline (10 mM Tris-HCl pH 7.5, 150 mM NaCl) containing 0.5% (v/v) Tween-20 (TBS-T) for 45 min at room temperature with gentle agitation. The α-VSVG antibody (V4888, Sigma) was added to the blocking buffer at a 1:5000 dilution for 1 h. The membrane was washed three times in TBS-T for 5 min each before incubation for 1 h with secondary antibody (anti-rabbit IgG HRP, 1:5000 dilution, 7074S, NEB) in TBS-T with 1% (w/v) skim milk powder. The membrane was then washed three times with TBS-T for 5 min each before developing using Clarity Max^TM^ Western ECL Substrate (1705062; BioRad) and imaged using a BioRad ChemiDoc XRS+.

### Error Prone PCR

Mutagenesis was adapted from Yang et al^12^. Four independent mutant plasmid libraries were constructed by mutagenizing *mraY* in plasmid pNG93 (P_lacUV5_::*mraY*) using Taq polymerase with Thermopol buffer (New England Biolabs, M0267L). The forward 5’- ACACTTTATGCTTCCGGCTC-3’ and reverse 5’- ACTGTTGGGAAGGGCGATCAAA-3’ primers were used to amplify *mraY* from pNG93. The resulting PCR products were purified using the Monarch® PCR & DNA Cleanup Kit (NEB, T1030) and used as “megaprimers” that are denatured and annealed to the original plasmid (pNG93) to amplify the vector backbone using Q5® High-Fidelity 2X Master Mix (NEB, M0492S). The reactions were then digested with DpnI to eliminate any remaining parental plasmid DNA. All four libraries were independently electroporated into NEB 10-beta electrocompetent cells (NEB, C3020K) and plated on LB agar supplemented with 15 μg/ml gentamicin at 37°C overnight.

Transformants were slurried in LB, and the resuspended cells were normalized to an OD_600_ = 10. Cells from 1 mL of resuspension were centrifuged and plasmid DNA was isolated from the cell suspension using the Monarch® Plasmid DNA Miniprep Kit (T1010). All four libraries were independently transformed into electrocompetent PA686 cells and plated on LBNS agar supplemented with 30 μg/ml Gent and grown overnight at 37°C. The resulting transformant colonies from each of the libraries were slurried in LBNS supplemented with 30 μg/ml Gent. Samples of each were normalized to OD_60_0 = 10 in LBNS + 10% (v/v) DMSO, and stored at −80°C. A sample from each library was then thawed and serial dilutions were plated on VBMM 30 μg/ml Gent with or without IPTG [50 μM], and grown at 30°C overnight. Individual colonies arising on the IPTG supplemented plates from each library were selected and re-streaked on VBMM with or without IPTG. Those that displayed IPTG dependence were further isolated, and the plasmids sent for sequencing. Clones identified to contain a single point mutation were further characterized. The mutated *mraY* genes were each amplified using Q5 High-fidelity polymerase (NEB) via colony PCR. The purified PCR product was digested with EcoRI and XmaI, and subsequently ligated into pPSV38 for validation of the suppression phenotype. All clones were sequence verified. MraY variants are listed in **Table S1**.

### Lipid II extraction

Cultures of PAO1, PA686 and MG1655 were grown at 37°C overnight, and MM119 at 30°C overnight. The next day, cultures were diluted to an OD_600_ of 0.01 and allowed to grow for 2 h at the above specified temperatures whereupon 1mM IPTG was added to induce expression of MraY. Cells were collected when the OD_600_ reached ∼0.5, and normalized to OD = 1 in a 1 mL volume. Pellets were collected by centrifugation at 21,000 x g and stored at -20°C until needed. Cells were resuspended in 1 mL LB and added to a mixture of 2:1 methanol : chloroform (3.5 mL total) in borosilicate glass tubes (16×100 mm, Fisher Scientific 1495935AA). Samples were vortexed for 1 min to form a single phase. Cell debris was collected by centrifugation for 10 min at 2000 × g, 21°C. The supernatant was transferred to a fresh borosilicate glass tube, and 2 mL of chloroform was added. The supernatant was acidified using 0.1N HCl to pH 1 as determined by pH indicator strips. The samples were vortexed for 1 min and centrifuged for 20 min at 2000 × g at 21°C to form a two-phase system. Using a glass pipette, as much of the aqueous upper layer was removed without disturbing the interface between the aqueous and organic phases and 1 mL methanol was subsequently added to form a single liquid phase upon vortexing. Samples were transferred to 1.5 mL Eppendorf tubes by glass pipette then dried by nitrogen stream at 40°C. Dried samples were dissolved in 150 μL of a mixture of methanol and chloroform (2:1) by vortexing then centrifuged at 21,000×g for 1 minute and dried by nitrogen stream at 40°C. This was repeated with 40 μL organic mixture, and finally crude lipid extracts were dissolved in 10 μL DMSO by vortexing. Extracts were stored at -20°C.

### Lipid II hydrolysis

Crude lipid II (LII) extracts were added (5 μL) to 5 μL of 0.2 M HCl, for a final concentration of 0.1 M HCl. Samples were boiled at 100°C for 15 min and then cooled to 4°C in a thermocycler. 10 μL of sodium borate pH 9 was added followed by 1 μL 0.5M NaOH to neutralize the solution. 2 μl of 100 mg/ml sodium borohydride was added and the samples were allowed to incubate for 30 min at room temperature. Following the incubation, 2 μl of 20% phosphoric acid was added to quench the reaction and the samples were mixed and immediately subjected to LC/MS analysis.

### LC/MS

High-resolution LC/MS traces of soluble LII hydrolysis products were obtained using the following protocol. Briefly, the hydrolyzed samples were subjected to LC/MS analysis (ESI, positive mode). A Waters Symmetry Shield RP8 column (3.5 µm, 4.6 mm X 150 mm) was used to separate hydrolysis products using the following gradient (A, H_2_O + 0.1% formic acid; B, acetonitrile + 0.1% formic acid; 0.5 ml/min): 0% B for 5 min, followed by a linear gradient of 0%-20% B over 40 min. Data was obtained on an Agilent 6546 LC-q-TOF Mass Spectrometer. Expected ion masses were extracted with a tolerance of 0.01 mass units.

### Purification of UDP-MurNAc pentapeptide

Accumulation of the precursor was performed as previously described^13^ with the following modifications. *Bacillus cereus* ATCC 14579 was grown in LB-lennox medium at 37°C until the OD_600_ reached between 0.7-0.8, at which point 130 μg/mL of chloramphenicol was added. After 15 minutes of incubation, 5 μg/ml of vancomycin was added and the cells allowed to incubate for another 60 min at 37°C with shaking. The culture was then cooled on ice and harvested by centrifugation (4000 x g, 20 min, SLC-6000 rotor, 4°C). Cells were collected and stored at -20°C until required.

Cells were resuspended in water (0.1 g wet weight/mL) and stirred into boiling water in a flask with stirring. Boiling was allowed to continue for another 15 minutes at which point the flask was removed from heat and allowed to cool to room temperature with stirring. After approximately 20 minutes the resuspension was cooled on ice and the debris was pelleted at 200,000 x g for 60 min at 4°C. The supernatant was removed and lyophilized. The lyophilized material was resuspended in water and acidified to pH 3 using formic acid (1mL/L culture extracted), centrifuged to remove precipitate, and immediately subjected to reversed phase high pressure liquid chromatography (RP-HPLC).

UDP-MurNAc pentapeptide was isolated by RP-HPLC on a Synergi 4u Hydro-RP 80A (250×10.0 mM). The column was eluted over a 30-min isocratic program (A, H_2_O + 0.1 % formic acid; B, acetonitrile + 0.1% formic acid; 4 ml/min), 4% B for 30 min at room temperature. The elution was monitored by UV at 254 nm. UDP-MurNAc-pentapeptide eluted approximately at 20 min in a single peak, which was verified by mass-spectrometry (1194.35 Da). Peak fractions were collected and lyophilized. The final product was resuspended in water for downstream use.

### Expression and Purification of PaMraY

For expression of *P. aeruginosa* MraY or MraY^T23P^, *E. coli* expression strain LSM9 containing pAM174 and the expression plasmid (pLSM116 or pLSM117) was grown in 1L TB supplemented with 2 mM MgCl_2_, kanamycin, and chloramphenicol at 37°C with shaking until the OD_600_ was 0.7. The cultures were cooled to 20 °C before inducing protein expression with 1mM IPTG and 0.1% (w/v) arabinose. Cells were harvested 19h post induction by centrifugation (6,000 x g, 15 min, 4°C). To purify FLAG-MraY or FLAG-MraY^T23P^, the cells were resuspended in lysis buffer B (50 mM HEPES pH 7.5, 150 mM NaCl, 20 mM MgCl_2_, 0.5 mM DTT) and lysed by passage through a cell disruptor (Constant systems) at 25 kpsi twice. Membranes were collected by ultracentrifugation (100,000 x g, 1h, 4°C). The membrane pellets were resuspended in solubilization buffer B (20 mM HEPES pH 7.0, 0.5M NaCl, 20% (v/v) glycerol, and 1% (w/v) DDM (Thermo Fisher)) and rotated end over end for 1h at 4°C before ultracentrifugation (100,000 x g, 1h, 4°C). The supernatant was supplemented with 2 mM CaCl_2_ and loaded onto a pre-equilibrated homemade M1 anti-FLAG antibody resin. The resin was washed with 25 column volumes (CVs) of wash buffer C (20 mM HEPES pH 7.0, 0.5M NaCl, 20% (v/v) glycerol, 2 mM CaCl_2_, 0.1% (w/v) DDM) and the bound protein was eluted from the column with five CVs of elution buffer (20 mM HEPES pH 7.0, 0.5M NaCl, 20% (v/v) glycerol, 0.1% (w/v) DDM, 5 mM EDTA pH 8.0, and 0.2 mg/mL FLAG peptide). Fractions containing the target protein were concentrated and the protein concentration was measured via the Bradford method. Proteins were aliquoted and stored at -80°C until required.

### MraY translocase in vitro assay

The assay was performed at 37°C in an assay buffer containing 20 mM HEPES pH 7.5, 500 mM NaCl, 20% (v/v) glycerol, and 0.1% (w/v) DDM, 10 mM MgCl_2_, 250 µM UDP-MurNAc pentapeptide, and 1.1 mM C55P (Larodan). Protein was added to initiate the reaction at a final concentration of 1.7 µM. At the appropriate time point the reaction was quenched by boiling for 3 min at 95°C. 1.5 units of alkaline phosphatase was added to the sample (NEB M0371L) and incubated at 25°C for 1h. The samples were heat quenched at 65°C to stop the reaction and were immediately loaded for analysis by LCMS. The samples were monitored by UV 254 and by MS (ESI, positive mode). A Thermo Fisher Hypersil Gold aQ C18 (150×4.6 mm 3 µm) HPLC column was used to separate the substrates and products using the following gradient program (A, H_2_O + 0.1% formic acid; B, acetonitrile + 0.1% formic acid; 0.4 ml/min): 4% B for 20 min. Data was obtained on an Agilent 6546 LC-q-TOF Mass Spectrometer.

### Preparation of lipopolysaccharide and immunoblotting

To isolate LPS from the *P. aeruginosa* strains containing the indicated plasmids, overnight cultures of each of the strains were allowed to grow in LB at 37°C containing 30 μg/mL Gent. The next day, cultures were diluted to an OD_600_ of 0.01 and allowed to grow at 37°C in 25 mL LB containing 30 μg/ml Gent. After 2 h, 1 mM IPTG was added and the cultures were allowed to grow for another 2h until cultures reached mid-log. 20 mL of culture was pelleted at 4000 x g for 12 min, and cells were resuspended in 1 mL LB and the OD_600_ was measured. The cells were pelleted again at 12,000 x g for 2 min, and resuspended in 1X LDS buffer (Invitrogen, NP00008) + 4% BME to an OD_600_ = 20. Samples were boiled at 95 °C for 10 min. Each sample was subjected to the NI Protein Assay (G Biosciences, 786-005) to determine the protein content in each sample. The lysates (50 μl) were then incubated at 55°C with 1.25 μl proteinase K (NEB, P8107S). After 1h of incubation, samples were boiled at 95°C for 10 min, and then frozen at -20°C until required.

Volumes of lysates corresponding to 20 μg of protein were then run on a Criterion XT 4- 12% Bis-Tris Precast Gel (Bio-Rad, 3450124) in MES running buffer (50 mM MES, 50 mM Tris base, 1 mM EDTA, 0.1% (w/v) SDS) for 1h 45 min at 100V constant. Glycan was transferred to nitrocellulose membranes as described above with the following differences: membranes were blocked for 1h at room temperature in 1% (w/v) skim milk, and were then incubated with anti-serotype O5 B-band at a 1:1000 dilution overnight at 4 °C (gift from L. Burrows). After three 15-mL TBST washes, membranes were incubated with anti-mouse HRP antibody (1:5000, NEB 7076S) for 1h at room temperature. Blots were developed as described above.

### Molecular dynamics simulations

For the coarse-grained MD, the structural model of the *E. coli* MraY dimer was aligned according to the plane of the membrane with memembed^14^, and then converted to the Martini 3 force field using the martinize protocol^15^. Bonds of 500 kJ mol^-1^ nm^-2^ were applied between all protein backbone beads within 1 nm. Proteins were built into 13 x 13 nm membranes composed of 40% POPE, and 10% each of POPG, CDL, lipid I, lipid II, C55-P, and C55-PP using the insane protocol^16^. Alternatively, membranes were built with 60% POPE, and 10% each of POPG, CDL, C55-P, and C55-PP. Lipid I, lipid II, C55-P and C55-PP parameters were from Orta et al. ^3^.Systems were solvated with Martini waters and Na^+^ and Cl^-^ ions to a neutral charge and 0.0375 M. Systems were minimized using the steepest descents method, followed by 1 ns equilibration using 5 fs time steps, then by 100 ns equilibration with 20 fs time steps, before 9 x 10 µs (complex membrane) or 5 x 10 µs (membrane without lipid I or lipid II) production simulations were run using 20 fs time steps, all in the NPT ensemble with the velocity-rescaling thermostat and semi-isotropic Parrinello-Rahman pressure coupling^17, 18^.

A pose of the *E. coli* MraY dimer with two lipid II molecules bound to the central cavity was selected for further analysis. All non-POPE lipids (except the two bound lipid II molecules) were deleted and the membrane allowed to shrink to 10 x 10 x 10.5 nm over 100 ns with positional restraints applied to the protein backbone. The resulting molecule was then converted to the atomistic CHARMM36m force field^19, 20^ using the CG2AT2 protocol^21^. Side chain pKas were assessed using propKa3.1^22^, and side chain side charge states were set to their default. Production simulations were run for 5 repeats of ca. 510 ns, using a 2 fs time step in the NPT ensemble with the velocity-rescale thermostat and semi-isotropic Parrinello-Rahman pressure coupling^17, 18^.

All simulations were run in Gromacs 2021.3^23^. Images were made in VMD^24^. Kinetic analysis of protein-lipid interactions and binding site identification were performed using PyLipID^25^. Density and contact analyses of atomistic MD simulations were performed using MDAnalysis^26, 27^. Contacts are defined as a distance of less than 4 Angstroms between Lipid II and MraY.

### Expression and purification of the YES complex

The YES complex was expressed as described previously^3^. Briefly, Δ*slyD* BL21(DE3) competent cells were transformed with pET22b-SlyD_1-154_ and pRSFDuet*Ec*MraY-E_ID21_ and plated in LB-agar containing 35 *µ*g/ml Kanamycin and 100 *µ*g/mL Ampicillin. The culture was grown in 2xYT media at 37^◦^C, 225 r.p.m., and induced at an OD_600_ of 0.9 with 0.4mM IPTG at 18^◦^C overnight. The culture was harvested by centrifugation for 10 minutes at 9,000*xg*, 4^◦^C followed by flash freezing.

The cells were lysed using a M-110L microfluidizer (Microfluidics) in 20mM Tris-HCl pH 7.5, 300 mM NaCl, 10% glycerol, 5mM *β*ME, 0.1mM PMSF, 0.1mM benzamidine. The lysate was cleared by a 20-minute centrifugation at 22,000*xg*. The membrane was isolated by ultracentrifugation at 167,424*xg* and solubilized in 10 mM HEPES pH 7.5, 300 mM NaCl, 5% Glycerol, 5mM *β*ME, 0.1mM PMSF, 0.1mM benzamidine, 10 mM imidazole and 1% dodecyl 4-O-*α*-D-glucopyranosyl-*β*-D-glucopyranoside (DDM). The extract was cleared by ultracentrifugation then nutated with 1mL NiNTA resin (Qiagen, Alameda, CA) at 4^◦^C for two hours. The resin was washed with five column volumes of wash buffer (10 mM HEPES pH 7.5, 150 mM NaCl, 5% glycerol, 5mM *β*ME, & 0.03% DDM) with 10mM imidazole and eluted in 20mL of wash buffer containing 200 mM imidazole. The eluent was further purified by SEC (Superdex 200 5/150 GL, Millipore Sigma) in 10mM HEPES pH 7.5, 75 mM NaCl, 5% glycerol, 5mM *β*ME and 0.03% DDM. Fractions were assessed by SDS-PAGE, concentrated, and directly used for cryo-EM sample preparation.

### Sample preparation for cryoEM

The protein sample was diluted to a concentration of 5mg/mL in 10mM HEPES pH 7.5, 75 mM NaCl, 2% glycerol, 5mM *β*ME, 0.03% DDM, and 1mM *E. coli* total lipid extract (Avanti Polar Lipids, 100600P). Quantifoil holey carbon films R1.2/1.3 300 Mesh, Copper (Quantifoil, Micro Tools GmbH) grids were glow discharged with a 2-minute 20Å plasma current using a Pelco easiGlow, Emitech K100X. Grids were prepared using a Vitrobot (FEI Vitrobot Mark v4 x2, Mark v3) by applying 3*µ*L of 5mg/mL YES(T23P) complex onto the grid followed by a 3.5 second blot using a +8-blot force and plunge frozen into liquid ethane.

### Data acquisition and analysis

Datasets were collected at a 105k magnification with a pixel size of 0.416 Å/pixel using a 300 kV cryo-TEM Krios microscope equipped with a Gatan K3 6k x 4k direct electron detector and a Gatan Energy Filter (slit width 20eV) in super-resolution mode using Serial EM. Movies with 40 frames were recorded with a total exposure dose of 60 e^-^/Å^2^ and a defocus range of -1.0 to -2.5 *µ*m. A total of 7,083 movies were gain reference and motion corrected using the patch motion correction built in function in cryosparc (v3.3.2) with a two-fold bin that resulted in a pixel size of 0.832 Å/pixel^28^. The contrast transfer function (CTF) was estimated using CTFFIND4^29^. A total of 3,885,223 particles were obtained by template picker using PDBID:8G01 as reference ^3^. Four ab-initio models were obtained using 500,000 particles, from which the best and worst volume were used to sort 4x binned particles through heterogeneous refinement.

Iterative rounds of heterogeneous and non-uniform refinement were performed before re-extracting particles using a 2x bin. This process was continued, and the resulting particles were re-extracted using a 1.3x bin. After several rounds of heterogeneous and non-uniform refinement, 575,243 particles were extracted without binning and used to create a map through non-uniform refinement. Using the MraY model from PDBID: 8G01, a mask covering only the density encompassing MraY was created using ChimeraX^30^. Density outside of this mask was removed using particle subtraction, followed by ab-initio modeling. The best fitting map was then used for further refinement using global CTF, heterogeneous, and non-uniform refinement. The final map with a 3.8Å resolution was composed by 287,765 particles and sharpened using the autosharpen module in PHENIX-1.19.2. The data collection, refinement and validation statistics can be found in Table S2.

## Notes

### Competing Interest Statement

The authors have declared no competing interest.

## REFERENCES

1. May, K. L. & Silhavy, T. J. Making a membrane on the other side of the wall. Biochimica Et Biophysica Acta Bba -Mol Cell Biology Lipids 1862, 1386–1393 (2017).

2. Silhavy, T. J., Kahne, D. & Walker, S. The Bacterial Cell Envelope. Csh Perspect Biol 2, a000414 (2010).

3. Huszczynski, S. M., Lam, J. S. & Khursigara, C. M. The Role of Pseudomonas aeruginosa Lipopolysaccharide in Bacterial Pathogenesis and Physiology. Pathogens 9, 6 (2019).

4. Rohs, P. D. A. & Bernhardt, T. G. Growth and Division of the Peptidoglycan Matrix. Annu. Rev. Microbiol. 75, 1–22 (2021).

5. Piepenbreier, H., Diehl, A. & Fritz, G. Minimal exposure of lipid II cycle intermediates triggers cell wall antibiotic resistance. Nat Commun 10, 2733 (2019).

6. Barreteau, H. et al. Quantitative high-performance liquid chromatography analysis of the pool levels of undecaprenyl phosphate and its derivatives in bacterial membranes. J Chromatogr B 877, 213–220 (2009).

7. Hartley, M. D. & Imperiali, B. At the membrane frontier: A prospectus on the remarkable evolutionary conservation of polyprenols and polyprenyl-phosphates. Arch Biochem Biophys 517, 83–97 (2012).

8. Jorgenson, M. A. & Young, K. D. Interrupting Biosynthesis of O Antigen or the Lipopolysaccharide Core Produces Morphological Defects in Escherichia coli by Sequestering Undecaprenyl Phosphate. J Bacteriol 198, 3070–3079 (2016).

9. Jorgenson, M. A., Kannan, S., Laubacher, M. E. & Young, K. D. Dead-end intermediates in the enterobacterial common antigen pathway induce morphological defects in Escherichia coli by competing for undecaprenyl phosphate. Mol Microbiol 100, 1–14 (2016).

10. D’Elia, M. A. et al. Lesions in teichoic acid biosynthesis in Staphylococcus aureus lead to a lethal gain of function in the otherwise dispensable pathway. J Bacteriol 188, 4183–9 (2006).

11. Lehrman, M. A. Biosynthesis of N -acetylglucosamine-P-P-dolichol, the committed step of asparagine-linked oligosaccharide assembly. Glycobiology 1, 553–562 (1991).

12. Sham, L.-T. et al. MurJ is the flippase of lipid-linked precursors for peptidoglycan biogenesis. Science 345, 220–222 (2014).

13. Sauvage, E., Kerff, F., Terrak, M., Ayala, J. A. & Charlier, P. The penicillin-binding proteins: structure and role in peptidoglycan biosynthesis. Fems Microbiol Rev 32, 234–258 (2008).

14. Zhao, H., Patel, V., Helmann, J. D. & Dörr, T. Don’t let sleeping dogmas lie: New views of peptidoglycan synthesis and its regulation. Mol Microbiol 106, 847–860 (2017).

15. Taguchi, A. et al. FtsW is a peptidoglycan polymerase that is functional only in complex with its cognate penicillin-binding protein. Nat Microbiol 4, 587–594 (2019).

16. Meeske, A. J. et al. SEDS proteins are a widespread family of bacterial cell wall polymerases. Nature 537, 634–638 (2016).

17. Rohs, P. D. A. et al. A central role for PBP2 in the activation of peptidoglycan polymerization by the bacterial cell elongation machinery. Plos Genet 14, e1007726 (2018).

18. Greene, N. G., Fumeaux, C. & Bernhardt, T. G. Conserved mechanism of cell-wall synthase regulation revealed by the identification of a new PBP activator in Pseudomonas aeruginosa. Proc National Acad Sci 115, 3150–3155 (2018).

19. Paradis-Bleau, C. et al. Lipoprotein Cofactors Located in the Outer Membrane Activate Bacterial Cell Wall Polymerases. Cell 143, 1110–1120 (2010).

20. Sardis, M. F., Bohrhunter, J. L., Greene, N. G. & Bernhardt, T. G. The LpoA activator is required to stimulate the peptidoglycan polymerase activity of its cognate cell wall synthase PBP1a. Proc National Acad Sci 118, e2108894118 (2021).

21. Typas, A. et al. Regulation of Peptidoglycan Synthesis by Outer-Membrane Proteins. Cell 143, 1097–1109 (2010).

22. Markovski, M. et al. Cofactor bypass variants reveal a conformational control mechanism governing cell wall polymerase activity. Proc National Acad Sci 113, 4788–4793 (2016).

23. Köhler, T., Donner, V. & Delden, C. van. Lipopolysaccharide as Shield and Receptor for R-Pyocin-Mediated Killing in Pseudomonas aeruginosa. J Bacteriol 192, 1921–1928 (2010).

24. Ge, P. et al. Action of a minimal contractile bactericidal nanomachine. Nature 580, 658–662 (2020).

25. Penterman, J. et al. Rapid Evolution of Culture-Impaired Bacteria during Adaptation to Biofilm Growth. Cell Reports 6, 293–300 (2014).

26. Chung, B. C. et al. Crystal Structure of MraY, an Essential Membrane Enzyme for Bacterial Cell Wall Synthesis. Science 341, 1012–1016 (2013).

27. Varadi, M. et al. AlphaFold Protein Structure Database: massively expanding the structural coverage of protein-sequence space with high-accuracy models. Nucleic Acids Res 50, D439–D444 (2022).

28. Jumper, J. et al. Highly accurate protein structure prediction with AlphaFold. Nature 596, 583–589 (2021).

29. Hakulinen, J. K. et al. MraY–antibiotic complex reveals details of tunicamycin mode of action. Nat Chem Biol 13, 265–267 (2017).

30. Oluwole, A. O. et al. Peptidoglycan biosynthesis is driven by lipid transfer along enzyme-substrate affinity gradients. Nat Commun 13, 2278 (2022).

31. Orta, A. K. et al. The mechanism of the phage-encoded protein antibiotic from ΦX174. Science 381, (2023).

## References

1. Studier, F. W. & Moffatt, B. A. Use of bacteriophage T7 RNA polymerase to direct selective high-level expression of cloned genes. J Mol Biol 189, 113–130 (1986).

2. Meeske, A. J. et al. SEDS proteins are a widespread family of bacterial cell wall polymerases. Nature 537, 634–638 (2016).

3. Orta, A. K. et al. The mechanism of the phage-encoded protein antibiotic from ΦX174. Science 381, (2023).

4. Guyer, M. S., Reed, R. R., Steitz, J. A. & Low, K. B. Identification of a Sex-factor-affinity Site in E. coli as gamma delta. Cold Spring Harb Symp Quant Biol 45, Pt 1:135–140 (1981).

5. Markovski, M. et al. Cofactor bypass variants reveal a conformational control mechanism governing cell wall polymerase activity. Proc National Acad Sci 113, 4788–4793 (2016).

6. Greene, N. G., Fumeaux, C. & Bernhardt, T. G. Conserved mechanism of cell-wall synthase regulation revealed by the identification of a new PBP activator in Pseudomonas aeruginosa. Proc National Acad Sci 115, 3150–3155 (2018).

7. Taguchi, A. et al. FtsW is a peptidoglycan polymerase that is functional only in complex with its cognate penicillin-binding protein. Nat Microbiol 4, 587–594 (2019).

8. Qiao, Y. et al. Detection of lipid-linked peptidoglycan precursors by exploiting an unexpected transpeptidase reaction. J Am Chem Soc 136, 14678–81 (2014).

9. Vvedenskaya, I. O. et al. Growth phase-dependent control of transcription start site selection and gene expression by nanoRNAs. Gene Dev 26, 1498–507 (2012).

10. Yunck, R., Cho, H. & Bernhardt, T. G. Identification of MltG as a potential terminase for peptidoglycan polymerization in bacteria: MltG lytic transglycosylase. Mol Microbiol 99, 700–718 (2015).

11. Choi, K.-H., Kumar, A. & Schweizer, H. P. A 10-min method for preparation of highly electrocompetent Pseudomonas aeruginosa cells: Application for DNA fragment transfer between chromosomes and plasmid transformation. J Microbiol Meth 64, 391–397 (2006).

12. Yang, D. C., Tan, K., Joachimiak, A. & Bernhardt, T. G. A conformational switch controls cell wall-remodelling enzymes required for bacterial cell division. Mol Microbiol 85, 768–781 (2012).

13. Kohlrausch, U. & Höltje, J. One-step purification procedure for UDP-N-acetylmuramyl-peptide murein precursors from Bacillus cereus. Fems Microbiol Lett 78, 253–257 (1991).

14. Nugent, T. & Jones, D. T. Membrane protein orientation and refinement using a knowledge-based statistical potential. Bmc Bioinformatics 14, 276 (2013).

15. Souza, P. C. T. et al. Martini 3: a general purpose force field for coarse-grained molecular dynamics. Nat Methods 18, 382–388 (2021).

16. Wassenaar, T. A., Ingólfsson, H. I., Böckmann, R. A., Tieleman, D. P. & Marrink, S. J. Computational Lipidomics with insane: A Versatile Tool for Generating Custom Membranes for Molecular Simulations. J Chem Theory Comput 11, 2144–2155 (2015).

17. Bussi, G., Donadio, D. & Parrinello, M. Canonical sampling through velocity rescaling. J Chem Phys 126, 014101 (2007).

18. Parrinello, M. & Rahman, A. Polymorphic transitions in single crystals: A new molecular dynamics method. J Appl Phys 52, 7182–7190 (1981).

19. Best, R. B. et al. Optimization of the Additive CHARMM All-Atom Protein Force Field Targeting Improved Sampling of the Backbone ϕ, ψ and Side-Chain χ1 and χ2 Dihedral Angles. J Chem Theory Comput 8, 3257–3273 (2012).

20. Huang, J. et al. CHARMM36m: an improved force field for folded and intrinsically disordered proteins. Nat Methods 14, 71–73 (2017).

21. Vickery, O. N. & Stansfeld, P. J. CG2AT2: an Enhanced Fragment-Based Approach for Serial Multi-scale Molecular Dynamics Simulations. J Chem Theory Comput 17, 6472–6482 (2021).

22. Søndergaard, C. R., Olsson, M. H. M., Rostkowski, M. & Jensen, J. H. Improved Treatment of Ligands and Coupling Effects in Empirical Calculation and Rationalization of pK a Values. J Chem Theory Comput 7, 2284–2295 (2011).

23. Berendsen, H. J. C., Spoel, D. van der & Drunen, R. van. GROMACS: A message-passing parallel molecular dynamics implementation. Comput Phys Commun 91, 43–56 (1995).

24. Humphrey, W., Dalke, A. & Schulten, K. VMD: Visual molecular dynamics. J Mol Graphics 14, 33–38 (1996).

25. Song, W. et al. PyLipID: A Python Package for Analysis of Protein–Lipid Interactions from Molecular Dynamics Simulations. J Chem Theory Comput 18, 1188–1201 (2022).

26. Gowers, R. et al. MDAnalysis: A Python Package for the Rapid Analysis of Molecular Dynamics Simulations. Proc. 15th Python Sci. Conf. 98–105 (2016) doi:10.25080/majora-629e541a-00e.

27. Michaud-Agrawal, N., Denning, E. J., Woolf, T. B. & Beckstein, O. MDAnalysis: A toolkit for the analysis of molecular dynamics simulations. J. Comput. Chem. 32, 2319–2327 (2011).

28. Punjani, A., Rubinstein, J. L., Fleet, D. J. & Brubaker, M. A. cryoSPARC: algorithms for rapid unsupervised cryo-EM structure determination. Nat Methods 14, 290–296 (2017).

29. Rohou, A. & Grigorieff, N. CTFFIND4: Fast and accurate defocus estimation from electron micrographs. J Struct Biol 192, 216–221 (2015).

30. Pettersen, E. F. et al. UCSF ChimeraX: Structure visualization for researchers, educators, and developers. Protein Sci 30, 70–82 (2021).

